# Glo1 reduction in mice results in age- and sex-dependent metabolic dysfunction

**DOI:** 10.1101/2025.01.24.634754

**Authors:** Ingrid Cely, Montgomery Blencowe, Le Shu, Graciel Diamante, In Sook Ahn, Guanglin Zhang, Jonnby LaGuardia, Ruoshui Liu, Zara Saleem, Susanna Wang, Richard Davis, Aldons J. Lusis, Xia Yang

**Affiliations:** Department of Integrative Biology and Physiology, University of California, Los Angeles, Los Angeles, CA, USA; Molecular Toxicology Interdepartmental Program, University of California Los Angeles, Los Angeles, CA, USA; Molecular, Cellular, and Integrative Physiology Interdepartmental Ph.D. Program, University of California, Los Angeles, Los Angeles, CA, United States of America; Departments of Medicine, Human Genetics, and Microbiology, Immunology, and Molecular Genetics, University of California, Los Angeles, Los Angeles, CA, USA; Department of Medicine, Division of Cardiology, David Geffen School of Medicine, University of California Los Angeles, Los Angeles, CA, USA; Department of Medicine, Human Genetics, and Microbiology, Immunology, and Molecular Genetics, University of California, Los Angeles, Los Angeles, CA, USA; Bioinformatics Interdepartmental Program, University of California, Los Angeles, Los Angeles, CA, USA; Institute for Quantitative and Computational Biosciences, University of California, Los Angeles, Los Angeles, CA, USA; Molecular Biology Institute, University of California, Los Angeles, Los Angeles, CA, USA

**Author notes:** **CORRESPONDENCE** Xia Yang, Ph.D. Department of Integrative Biology and Physiology University of California, Los Angeles, Los Angeles, CA 90095, USA.

**Keywords:** Glyoxalase 1, Sex difference, Advanced glycation endproducts, Metabolic syndrome, Obesity, Glucose dysregulation

## Abstract

**Objectives:** Advanced glycation end products (AGEs) have been implicated as an important mediator of metabolic disorders including obesity, insulin resistance, and coronary artery disease. Glyoxalase 1 (Glo1) is a critical enzyme in the clearance of toxic dicarbonyl such as methylglyoxal, precursors of AGEs. The role of AGE-independent mechanisms that underly Glo1-induced metabolic disorders have yet to be elucidated.

**Methods:** We performed a longitudinal study of female and male *Glo1* heterozygous knockdown (*Glo1^+/-^*) mice with ∼50% gene expression and screened metabolic phenotypes such as body weight, adiposity, glycemic control and plasma lipids. We also evaluated atherosclerotic burden, AGE levels, and gene expression profiles across cardiometabolic tissues (liver, adipose, muscle, kidney and aorta) to identify pathway perturbations and potential regulatory genes of Glo1 actions.

**Results:** Partial loss of *Glo1* resulted in obesity, hyperglycemia, dyslipidemia, and alterations in lipid metabolism in metabolic tissues in an age- and sex-dependent manner. *Glo1^+/-^* females displayed altered glycemic control and increased plasma triglycerides, which aligned with significant perturbations in genes involved in adipogenesis, PPARg, insulin signaling, and fatty acid metabolism pathways in liver and adipose tissues. Conversely, *Glo1^+/-^* males developed increased skeletal muscle mass and visceral adipose depots along with changes in lipid metabolism pathways. For both cohorts, most phenotypes manifested after 14 weeks of age. Evaluation of methylglyoxal-derived AGEs demonstrated changes in only male skeletal muscle but not in female tissues, which cannot explain the broad metabolic changes observed in *Glo1^+/-^*mice. Transcriptional profiles suggest that altered glucose and lipid metabolism may be partially explained by alternative detoxification of methylglyoxal to metabolites such as pyruvate. Moreover, transcription factor (TF) analysis of the tissue-specific gene expression data identified TFs involved in cardiometabolic diseases such as Hnf4a (all tissues) and Arntl (aorta, liver, and kidney) which are female-biased regulators and whose targets are altered in response to *Glo1^+/-^*.

**Conclusions:** Our results indicate that Glo1 reduction perturbs metabolic health and metabolic pathways in a sex- and age-dependent manner without significant changes in AGEs across metabolic tissues. Rather, tissue-specific gene expression analysis suggests that key transcription factors such as Hfn4a and Arntl as well as metabolite changes from alternative methylglyoxal detoxification such as pyruvate, likely contribute to metabolic dysregulation in *Glo1^+/-^* mice.

## 1. INTRODUCTION

There is a global epidemic of obesity and related metabolic disorders such as cardiovascular disease (CVD) and type 2 diabetes (T2D). Despite current preventative and therapeutic strategies, prevalence remains on the rise and improved mapping of disease mechanisms may offer new biological insights and therapeutic targets. Glyoxalase 1 (Glo1) has been linked to obesity[1] [1, 2], glycemic control [3], insulin sensitivity [4], aortic endothelial cell dysfunction [5], non-alcoholic fatty liver disease (NAFLD) [6], skeletal muscle dysfunction [7] and CVD [8], suggesting its importance in regulating metabolic health [9, 10]. Although AGEs have been the center focus of previous studies to explain Glo1-mediated dysfunction, AGE-independent mechanisms remain underexplored. Glo1 is part of the glyoxalase system, which is comprised of ubiquitous enzymes Glo1, Glyoxalase 2 (Glo2), and a catalytic amount of reduced glutathione (GSH) [11]. The glyoxalase system serves as a defense mechanism against the accumulation of reactive dicarbonyls such as methylglyoxal (MG), a cytotoxic byproduct of glycolysis [12]. Once generated, MG and GSH can spontaneously form a hemithioacetal, the substrate for Glo1. The enzyme-substrate complex can then generate S-D-Lactoylglutathione, which is hydrolyzed by Glo2 to produce the final products, D-lactate and GSH. In essence, Glo1 is the main enzyme that initiates the process of converting a reactive metabolite such as MG into D-lactate, a non-toxic and more stable product.

The impairment of the glyoxalase system can lead to the accumulation of MG, resulting in dicarbonyl stress. Accumulated MG can react with DNA, lipids and amino acid residues of proteins. Specifically, MG readily reacts with arginine and lysine residues to irreversibly generate advanced glycation end products (AGEs), making MG a major precursor for AGEs. The two most abundant MG-derived AGEs are *N-*(5-hydro-5-methyl-4-imidazolon-2-yl)-ornithine (MGH1) and carboxyethyl-lysine (CEL). The interaction between AGEs and the receptor for advanced glycation endproducts (RAGE), plays a major role in the pathology of various diseases, including multiple cancers, Alzheimer’s Disease, NAFLD, insulin resistance, diabetes and its complications including micro- and macrovascular disease, retinopathy, nephropathy and neuropathy [4, 13–16]. With a compromised glyoxalase system, an alternative detoxification defense system against MG can partially compensate to mitigate the effects caused by dicarbonyl stress and AGEs. This defense system includes enzymes, aldehyde dehydrogenase (ALDH) and aldoketo reductase (AKR), that act to metabolize MG to pyruvate and hydroxyacetone, respectively [17].

Despite the known role of the MG-AGE pathway in obesity and metabolic dysfunction, previous studies in fish and mice have revealed that full Glo1 knockouts, does not always result in elevated MG, AGEs or even an observable phenotype, likely through compensation of ALDH and AKR [18]. On the other hand, Glo1 knockouts in drosophila and *C.elegans* have demonstrated elevated MG and AGEs [19] [20]. Collectively, these highlight the variability of underlying mechanisms at play in different Glo1 model organisms and species. Although Glo1 knockout studies have identified a role for MG-AGE leading to the development of obesity and metabolic dysfunction, the human population rarely exhibits a complete loss of Glo1 activity with an incomplete knowledge of the mechanisms behind Glo1 function [21]. To explore a more physiologically relevant state, we have opted to study the effect of Glo1 reduction using the Glo1*^+/-^*mouse on the C57BL/6 background, which exhibits a 45-65% reduction in Glo1 enzymatic activity. We not only examined the mechanisms underlying the Glo1 effect on obesity and metabolic dysfunction, but also determined the impact of sex and age over a six-month period.

Our results showed that compared to male *Glo1^+/-^* mice, female *Glo1^+/-^* mice manifested more severe phenotypes including elevated glucose and impaired lipid metabolism. However, AGEs were only increased in male muscle, which do not explain the female-biased phenotypes. Gene expression, metabolic pathway, and network analyses support the involvement of female-biased transcription factors such as HNF4A as well as metabolite products such as pyruvate from detoxification of MG in the metabolic reprogramming in females. Furthermore, integration of the tissue-specific transcriptomic signatures altered in female *Glo1^+/-^* mice with human genome-wide association studies (GWAS) provided additional mechanistic and clinical insights into the connection between *Glo1* and multiple human metabolic traits and diseases. Our findings support an important age- and sex-dependent role for *Glo1* in mediating obesity and associated comorbidities through alterations of numerous metabolic pathways without major changes in AGEs.

## 2. METHODS

### 2.1 Heterozygous *Glo1* knockdown mice

The *Glo1^+/-^* mice were originally generated in Dr. Michael Brownlee’s lab by injecting Glo1-targeting shRNA lentivirus into C57/B6 mouse embryos. Mice whose genome contained a single copy of the insert were identified by Southern blotting and used to establish founder lines. Glo1 mRNA and protein levels were determined by quantitative PCR and Western blot analysis and further confirmed by measurement of Glo1 activity. Heterozygous offspring of the founder had a 45–65% decrease in tissue Glo1 activity, and these mice were used in all experiments [22]. *Glo1^+/-^*embryos were received as a gift from Dr. Abraham Palmer’s lab and re-derived at the University of California, Los Angeles (UCLA) by Division of Laboratory Animal Medicine (DLAM) the implantation of the embryo into a surrogate C57BL/6 female mouse. In our lab, offspring male *Glo1^+/-^*mice were mated with female wildtype mice to expand the mouse colony. Genotype of all mice were confirmed by PCR as detailed below. All experiments on *Glo1^+/-^* mice and littermate controls were conducted in accordance with the United States National Institutes of Health Guide for the Care and Use of Laboratory Animals and were approved by the UCLA Animal Research Committee.

### 2.2 Genotyping and husbandry

Genomic DNA was extracted from mouse ears (n=10-13 per sex) and analyzed by PCR using Kapa Mouse Genotyping Kit (Kapa Biosystems, Wilmington, MA, USA). Glo1 knockdown primers were as follows: forward (5’-GCTTCTCCCACAAGTCTGTG-3’) and reverse (5’-GGTACAGTGCAGGGGAAAGA-3’). Gapdh primers served as the control and were as follows: forward (5’-AACTTTGGCATTGTGGAAGG-3’) and reverse (5’-ACACATTGGGGGTAGGAACA-3’). Mice were provided with *ad libitum* standard chow diet (Newco Distributors Inc., Rancho Cucamonga, CA, USA) and water. Food and water intake were measured weekly to monitor caloric intake between groups.

### 2.3 Glo1 enzyme activity assay

Protein samples were prepared from 50 mg of liver and kidney tissues and used to determine Glo1 enzyme activity according to manufacturer’s instructions (Sigma-Aldrich, St. Louis, MO, USA). Briefly, protein extraction of tissues was mixed with substrate and cosubstrate and incubated for 20 minutes. Samples were precipitated with perchloric acid on ice, vortexed and then centrifuged at 14,000 rpm for 5 minutes. Supernatant was collected then transferred to a 96 UV well plate in duplicates and absorbance was read at 240nm using a microplate reader (Biotek, USA). A blank was also prepared and processed as previously mentioned without the addition of the protein. Glo1 activity was determined by measuring the production of S-Lactoylglutathione. One unit of Glo1 is the amount of enzyme that will convert 1.0 μmol of S-Lactoylglutathione from methylglyoxal and reduced glutathione per minute at pH 6.6 and *25 °C*. Once the absorbance was determined from each sample, a calculation provided on the manufacturer’s instructions was used to obtain the final Glo1 activity in units/L.

### 2.4 Body weight and composition

For both *Glo1^+/-^* and littermate control mice (n=10-13/genotype/sex), starting at 3-weeks of age, body weight was monitored weekly until the age of 28 weeks. Starting at 5-weeks of age, fat mass and lean muscle was monitored biweekly until the age of 27 weeks. Body weight was obtained by weighing mice on a scale and fat and lean mass data was obtained using nuclear magnetic resonance (NMR) (Brucker, Madison, WI, USA). Statistical analysis was done by 2-way ANOVA with repeated measures, followed by Bonferroni correction.

### 2.5 Intraperitoneal glucose tolerance test (IPGTT)

Mice underwent a glucose tolerance test at several timepoints of the experiment (5, 12, 23 and 33 weeks old). Mice were fasted overnight for 14 hours prior to IPGTT. A 20% glucose solution (Sigma-Aldrich, St. Louis, MO, USA) was prepared for intraperitoneal (IP) injection (2g of glucose per kg body mass). Blood samples (<5uL) for glucometer reading were obtained via mouse-tail incision. Blood glucose was measured at 15-, 30-, 60- and 120-minutes post glucose injection to test clearance of glucose.

### 2.6 Plasma lipid, glucose and insulin quantification

Quantification of blood plasma lipids, glucose and insulin was conducted at several timepoints of the experiment (7, 12 and 28 weeks of age). Prior to blood collection mice were fasted overnight for 14 hours. Fasted mice were anesthetized using isoflurane prior to blood collection by retro-orbital bleeding using a microcapillary tube. The collected blood was put into K2 EDTA tubes and placed on ice. The amount of blood obtained was 1% of the total body weight. Blood was then centrifuged at 1500g for 10 minutes and plasma was collected and stored in -80°C. Plasma triglycerides (TG), total cholesterol (TC), unesterfied cholesterol (UC), free fatty acids (FFA), high-density lipoprotein (HDL) cholesterol, low-density lipoprotein (LDL) cholesterol, glucose and insulin were analyzed by enzymatic colorimetric assays as previously described [23]. Very low-density lipoprotein (VLDL) cholesterol was calculated using the formula: VLDL= (TG/5).

### 2.7 Characterization of atherosclerotic lesions

Mouse hearts were mounted with optimal cutting temperature (OCT) compound (VWR, Radnor, PA, USA) and placed in -80°C for histological purposes. Using a stereomicroscope, the heart was sliced longitudinally into 10 µm thick sections and fixed in 80% 2-propanol. The aortic sinus was stained with Oil Red O (Abcam, Cambridge, MA, USA) and lesions were quantified. Sections were fixed in 10% formalin for 10 minutes, rinsed 3 times in 1X PBS, placed in Oil Red O solution for 10 minutes, rinsed in tap water and counterstained with hematoxylin for 1 minute. An inverted phase-contrast microscope (Eclipse TE 300; Nikon Co., Tokyo, Japan) was used for identification of lesions.

### 2.8 Total RNA extraction, cDNA synthesis and qPCR

Mouse liver, gonadal adipose and skeletal muscle tissues were flash frozen in liquid nitrogen immediately after euthanasia. RNA extraction was performed using RNeasy Mini Kit (Qiagen, Germantown, MD, USA) following the manufacturer’s instructions. RNA concentration was determined by Nanodrop ND-1000 Spectrophotometer (Thermo Fisher Scientific, Waltman, MA, USA). cDNA synthesis was performed using Applied Biosystems High-Capacity cDNA Reverse Transcription Kit (Thermo Fisher Scientific, Waltman, MA, USA). Expression of *Ager, Akr1a1, Aldh1a1, Lipin1, Acc1, Fasn, Elovl6, Scd1, Srebp1c, Dgat1 and Dgat2* was analyzed by qPCR using the Applied Biosystem Real-Time PCR Instrument (Thermo Fisher Scientific, Waltman, MA, USA) and SYBR Green Master Mix (Thermo Fisher Scientific, Waltman, MA, USA). Full primer sequences are provided in Supplementary Table 1.

### 2.9 Protein extraction from metabolic tissues

In preparation of ELISA and Western Blot experiments, protein extraction from liver, gonadal adipose and skeletal muscle was performed according to the Abcam ELISA sample preparation guide on frozen tissue as follows: Approximately 50 mg of frozen liver and skeletal muscle tissue and 100 mg of frozen gonadal adipose tissue were homogenized in tissue extraction buffer consisting of 1% Triton X-100 (Thermo Fisher Scientific, Hudson, NH, USA), 100 mM Tris (pH 7.4), 150 mM NaCl, 1 mM EGTA, 1 mM EDTA, 0.5% sodium deoxycholate and protease and phosphatase inhibitor cocktail (Abcam, Cambridge, MA, USA). The homogenized tissue was then centrifuged at 13000 rpm for 20 minutes at 4°C and the supernatant containing the soluble protein extract was collected. Total protein concentration was evaluated by Bicinchoninic Acid (BCA) Protein Assay (Pierce, Rockford, IL, USA). Lysate was diluted to 2000µg/mL total protein for liver and skeletal muscle tissues and 500µg/mL total protein for gonadal adipose prior to running ELISA. In preparation of Western Blot experiments, protein extraction from liver and kidney was performed using approximately 100 mg of frozen tissue as described above. Lysate was diluted to 80µg prior to running Western Blot experiment.

### 2.10 ELISA quantification of AGEs

The two main AGEs MGH1 and CEL were quantified using the protein extracts from liver, gonadal adipose and skeletal muscle tissues. Quantification of MGH1 was performed using OxiSelect™ Methylglyoxal (MG) Competitive ELISA Kit (Cell Biolabs Inc., San Diego, CA, USA). Quantification of CEL was performed using OxiSelect™ N-epsilon-(Carboxyethyl) Lysine (CEL) Competitive ELISA (Cell Biolabs, San Diego, CA, USA). ELISA experiments were analyzed using a microplate reader (Biorad, Hercules, CA, USA) and Gen5 data analysis software.

### 2.11 Western Blotting for Glo1 and Rage quantification

Using liver and kidney tissues, 80µg of protein lysate was loaded on the SDS-PAGE gel according to manufacturer’s instruction (General Protocol for Western Blotting; BioRad). The primary antibody was incubated overnight at 4◦C. Following this, HPR-conjugate antibody was incubated for 1 hour in room temperature. Lastly, signals were captured using the Super Signal West Pico PLUS Chemiluminescent kit (Thermo Scientific REF34580) and visualized on Bio Rad imager using the Image Lab 3.01 software.

### 2.12 Total RNA isolation, microarray profiling, and the identification of differentially expressed genes (DEGs)

Liver, gonadal adipose, kidney and aorta from 34-week-old female mice were flash frozen in liquid nitrogen. About 10-15 mg tissue for liver, kidney and aorta and 30 mg for gonadal adipose tissue was homogenized and processed using RNeasy mini kit (QIAGEN GmbH, Hilden, Germany) for RNA isolation. RNA concentration was determined using Nanodrop (Thermo Fisher Scientific, MA, USA) and Bioanalyzer (Agilent Technologies, CA, USA). A total of 23 samples passing quality control (RIN > 7.0) were sent to the UCLA Neuroscience Genomics Core facility for labeling and hybridization using the Illumina MouseRef-8 v2.0 array. There were 11 samples from C57/B6 WT control mice (3 samples/tissue, with the exception of aorta which had 2 samples), and 12 samples from *Glo1^+/-^* mice (3 samples/tissue). The gene expression data were deposited in Gene Expression Omnibus with accession number GSE118034. The Illumina array data were analyzed using the lumi Bioconductor package within R [24]. Variance stabilization transformation was applied, followed by robust spline normalization. Differentially expressed genes (DEGs) between groups were identified using a linear model. False discovery rates (FDR) for the differential expression p-values were determined using the q-value Bioconductor package [25]. FDR (q-value) < 0.1 were used as cutoffs to determine significant DEGS for downstream pathway, and suggestive DEGs with p < 0.05 were used in integrative genomics analyses.

### 2.13 Functional annotation of *Glo1^+**/**^* DEGs

*Glo1^+/-^* DEGs identified from microarray analysis were annotated for their potential biological functions using canonical pathways or functional categories from various databases including Kyoto Encyclopedia of Genes and Genomes (KEGG) [26], BioCarta (http://www.biocarta.com/genes/index.asp), Reactome [27], and gene sets derived from Gene Ontology [28]. Fisher’s exact test was performed to calculate the enrichment p-values for each pathway or functional category within the up- and down-regulated DEGs followed by multiple testing correction using Storey’s method [22]. In addition, we also utilized gene set enrichment analysis (GSEA) to highlight tissue specific biological pathways as well as the distibution of up and downregulated genes within those pathways.

### 2.14 Identification of enriched transcription factor downstream targets among Glo1*^+/-^*DEGs

To identify potential upstream regulators of *Glo1^+/-^*DEGs, we assessed whether downstream target genes of transcription factors (TFs) were enriched in *Glo1^+/-^* DEGs using the binding analysis for regulation of transcription (BART) computational tool. We used an Irwin-Hall p-value cutoff (p<0.01) to identify TFs.

### 2.15 Assessing enrichment of *Glo1^+/-^* DEGs for genes mapped to genetic loci of human metabolic traits

To test whether there is a link between the genes identified from our study and human metabolic traits, we curated a total of 15 full sets of GWAS summary statistics in the public data repositories (i.e., disease association p-values of all the tested genetic risk variants in the form of single nucleotide polymorphisms, or SNPs) in existing large-scale GWAS for metabolic traits, including obesity (adult BMI [29], adult hip circumference [30], adult waist circumference [30], adult waist-hip ratio [30], T2D [31], CAD [32], glucose homeostasis traits HbA1c [33], fasting glucose [34], fasting insulin [34], the homeostatic model assessment for beta-cell function (HOMA-B) [35], insulin resistance (HOMA-IR) [35], and lipid profiles (TC, TG, LDL, HDL) [36]. For each GWAS, we removed SNPs with a minor allele frequency < 0.05. For SNPs that are in linkage disequilibrium with r^2^ > 0.5, only the SNP with the strongest disease association was kept. We then used Marker Set Enrichment Analysis (MSEA) to determine whether the genes affected in our *Glo1^+/-^* animal model were also enriched for SNPs that demonstrated evidence for disease association in humans. MSEA assesses whether a defined group of genes (in this study, DEGs) is enriched for disease-associated SNPs compared to random chance [37]. GWAS reported SNPs within the 50kb up/down-stream of a gene was mapped to the corresponding gene. For the list of SNPs mapped to each gene-set, MSEA tested the enrichment of SNPs for disease association using a chi-like statistic. The null background was estimated by permuting gene labels to generate random gene sets matching the gene number of each DEG set, while preserving the assignment of SNPs to genes. For each DEG set, 10000 permuted gene sets were generated and enrichment p-values were determined from a Gaussian distribution approximated using the enrichment statistics from the 10000 permutations and the statistics of the real DEG set. Finally, Benjamini-Hochberg FDR was estimated across all DEG sets tested for each GWAS.

## 3. RESULTS

### 3.1 Reduction of *Glo1* gene expression and enzymatic activities in *Glo1^+/-^* mice

To confirm genotype of mice and establish experimental groups, all mice were first genotyped by PCR at 3 weeks of age, as previously mentioned. PCR results showed that Glo1 band was absent in *Glo1^+/-^*mice (Supplementary Figure 1A). At 28-weeks of age, western blot analysis showed that protein expression of Glo1 in liver and kidney tissue was reduced in *Glo1^+/-^* mice compared to wildtype controls. In addition, Glo1 enzyme activity was assessed and results showed that Glo1 activity trended lower in *Glo1^+/-^* mice compared to wildtype controls at 28 weeks in liver and kidney tissues (p=0.0675; Supplementary Figure 1B). Lastly, expression profiling by microarray showed that *Glo1* expression was lowered by 46% in adipose (p = 3.2e-3), 55% in aorta (p = 1.3e-3), and 46% in liver (p = 1.5e-4), confirming the Glo1 reduction mice at 34 weeks (Figure 5D).

### 3.2. *Glo1* reduction affects body composition

To investigate the effects of *Glo1^+/-^* and determine when phenotypes would manifest, body weight was monitored weekly starting at weaning age (3 weeks) until 28 weeks of age (n=10-13/group) (Figure 1A,1B). Fat and muscle mass were monitored biweekly starting at 5 weeks of age until 27 weeks of age. Compared to WT counterparts, female *Glo1^+/-^*mice started exhibiting significantly increased body weight at 17 weeks and male *Glo1^+/-^*mice at 15 weeks and this trend continued until 28 weeks of age (Figure 1B; 2-way ANOVA with repeated measures). Female and male *Glo1^+/-^* mice both showed significantly increased fat mass at 15 weeks and continued through the end of the experiment at 28 weeks (Figure 1C). Interestingly, male *Glo1^+/-^*mice showed significantly increased muscle mass at 11 weeks of age until 28 weeks compared to control mice, whereas no change in muscle mass was observed between female groups (Figure 1D). Lastly, both female and male *Glo1^+/-^* mice demonstrated significantly increased adiposity (Figure 1E) from 15-28 weeks compared to their corresponding controls. Food and water intake was also concurrently monitored and these finding show that there was no consistent difference in water or food intake between groups for a 27-week duration period to account for significant differences in body composition in mice (Supplementary Figure 2A-2D).

**Figure 1.**
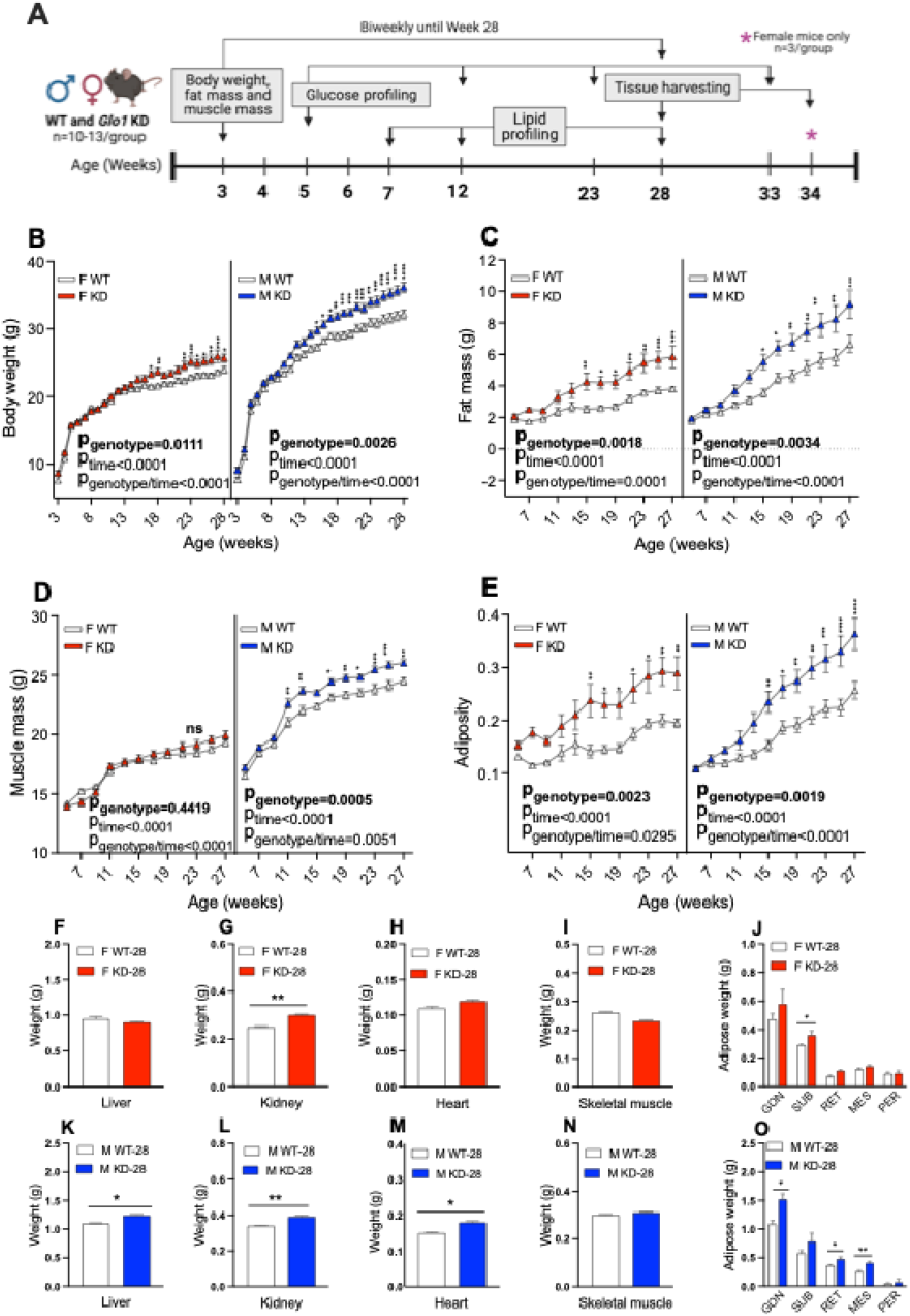
Characterization of body composition phenotypes in *Glo1^+/-^* mice. **(A)** Overview of experiment design with timepoints shown to indicate data collection in mice. Body weight for **(B)** female (n=12-13/group) and male (n=11-14/group) mice were monitored weekly starting from weaning (3 weeks) until 28 weeks of age. **(C)** Fat mass and **(D)** muscle mass was monitored biweekly starting from age 5 weeks of age until 27 weeks of age. **(E)** Adiposity (fat to muscle mass ratio) was monitored biweekly starting from age 5 weeks until 27 weeks. Female **(F)** liver, **(G)** kidney, **(H)** heart and **(I)** bilateral skeletal muscle tissue was weighed at 28 weeks of age. **(J)** Female adipose tissues including gonadal, subcutaneous, retroperitoneal, mesenteric and perirenal adipose tissues were weighed at 28 weeks of age. Male **(K)** liver, **(L)** kidney, (**M)** heart and **(N)** bilateral skeletal muscle tissue was weighed at 28 weeks of age. **(O)** Male adipose tissues including gonadal, subcutaneous, retroperitoneal, mesenteric and perirenal adipose tissue were weighed at 28 weeks of age. Data is shown as mean ± SEM and analyzed using 2-way ANOVA with repeated measures to obtain the statistical significance of the main effects of genotype and time as well the interaction between genotype and time. For individual time points, the difference between genotypes is assessed using the post-hoc Bonferroni’s multiple comparisons corrections following 2-way ANOVA with repeated measures. *p<0.05, **p<0.01, ***p<0.001.

Metabolic tissue weights including liver, kidney, heart, skeletal muscle, and fat pads (gonadal, subcutaneous, retroperitoneal, mesenteric and perirenal adipose) were also compared between groups at 28 weeks of age (Figure 1F-1O). Female *Glo1^+/-^* kidneys and subcutaneous adipose were significantly increased compared to controls (Figure 1F-1J). Male *Glo1^+/-^*livers, kidneys, hearts, gonadal, retroperitoneal and mesenteric adipose tissues was significantly increased compared to controls (Figure 1K-1O). Male *Glo1^+/-^* lean mass, determined by NMR, was significantly increased compared to controls, while the vastus lateralis muscle dissected from the bilateral hindlegs, was not. Together, these results suggest that decreased *Glo1* expression results in obesity post-developmentally in both female and male in mice.

### 3.3 *Glo1* reduction induces glucose intolerance in female but not male mice

To assess the impact of *Glo1* reduction on glucose tolerance over time, we performed IPGTT experiments on 14-hour fasted mice at 7, 12, 23, 28 and 33 weeks of age (Figure 2). Additional mice for both sexes in each group (n=5/group) were exempt from the 28-week endpoint to further assess glycemic phenotypes at the 33-week endpoint (Figure 2D, 2H). We observed significantly increased glucose AUC in female *Glo1^+/-^*mice at 23 weeks (n=8/group) and 33 weeks (n=5/group) compared to WT (Figure 2I) but no differences in male *Glo1^+/-^* mice (Figure 2L). Moreover, female *Glo1^+/-^* showed no difference between genotypes when comparing fasting blood glucose levels (Figure 2J) or insulin levels (Figure 2K), but male *Glo1^+/-^* had significantly decreased glucose levels at 12 weeks of age (Figure 2M) without any differences observed in insulin levels (Figure 2N). This data demonstrates that *Glo1* reduction results in the development of glucose intolerance in female but not male mice.

**Figure 2.**
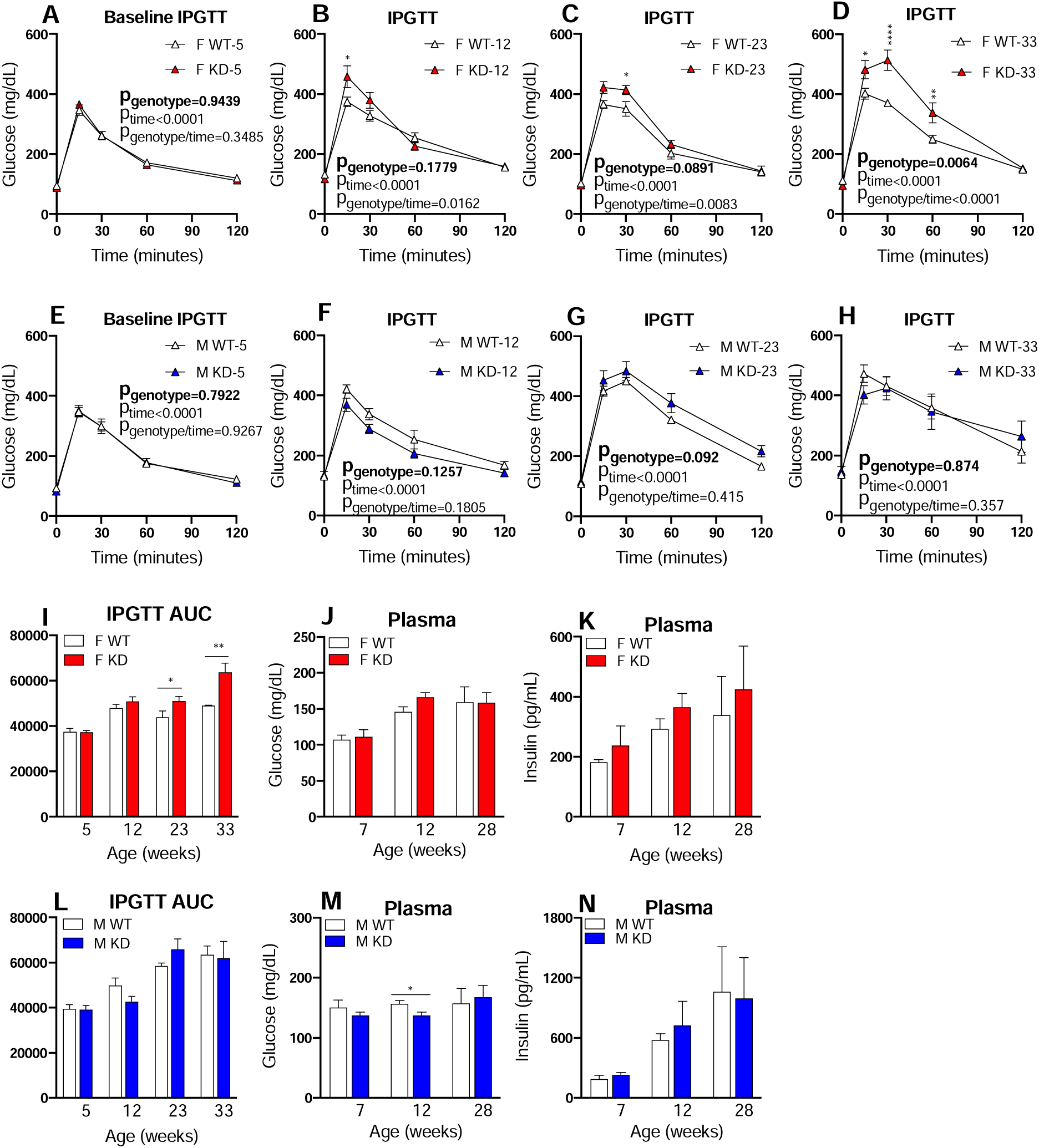
Characterization of glucose metabolism phenotypes in *Glo1^+/-^* mice. **(A)** Baseline intraperitoneal glucose tolerance test (IPGTT) was performed on fasted female mice at 5 weeks of age (n=13). Follow up IPGTT experiments were performed on female mice at **(B)** 12 (n=13), **(C)** 23 (n=13) and **(D)** 33 weeks of age (n=5). **(I)** Quantification for area under the curve (AUC) was determined for female mice for all times points. **(E)** Baseline IPGTT was performed on fasted male mice at 5 weeks of age (n=13). Follow up IPGTT experiments were performed on male mice at **(F)** 12 (n=13), **(G)** 23 (n=13) and **(H)** 33 weeks of age (n=5). **(L)** Quantification for area under the curve (AUC) was determined for male mice for all times points. Blood plasma **(J)** glucose and **(K)** insulin levels were quantified in fasted female mice at 7 (n=9), 12 (n=9) and 28 weeks of age (n=9). Blood plasma **(M)** glucose and **(N)** insulin levels were quantified in fasted male mice at 7 (n=9), 12 (n=9) and 28 weeks of age (n=9). IPGTT data was analyzed using 2-way ANOVA followed by Bonferroni’s multiple comparisons tests to obtain the statistical significance of the main effects of genotype and time as well the interaction between genotype and time. AUC data was analyzed using 1-way ANOVA followed by Bonferroni’s multiple comparisons tests. *p<0.05, **p<0.01, ***p<0.001. Plasma glucose and insulin data is presented as mean ± SEM and statistical significance among groups was calculated by Student t-test. *p<0.05, **p<0.01, ***p<0.001.

### 3.4 Glo1 reduction results in dyslipidemia in a sex-dependent manner

To assess the impact of *Glo1* reduction on lipid phenotypes over time, we measured plasma TG, TC, UC, FFA, HDL, LDL and VLDL levels after a 14 hour overnight fast (Figure 3A-3N). Hyperlipidemia, characterized by significantly elevated TG (Figure 3A) and VLDL (Figure 3G) levels, was observed in female *Glo1^+/-^* mice at 28 weeks compared to WT counterparts. Surprisingly, male *Glo1^+/-^*mice revealed hypolipidemia, a contrasting lipid phenotype to female *Glo1^+/-^* mice, characterized by significantly decreased TG (Figure 3H), TC (Figure 3I) and HDL (Figure 3L) at 28 weeks without changes in other lipids. While both female and male *Glo1^+/-^*mice show evidence for dyslipidemia at 28 weeks, the metabolic phenotypes manifested are sex-dependent. Overall, female *Glo1^+/-^* mice appear to reveal greater susceptibility to increases in circulating lipids compared to WT mice, while male *Glo1^+/-^* mice demonstrate the opposite trend and show decreases in circulating lipids compared to control groups. These findings provide evidence that *Glo1* reduction affects certain plasma lipids in both a sex- and age-dependent manner.

**Figure 3.**
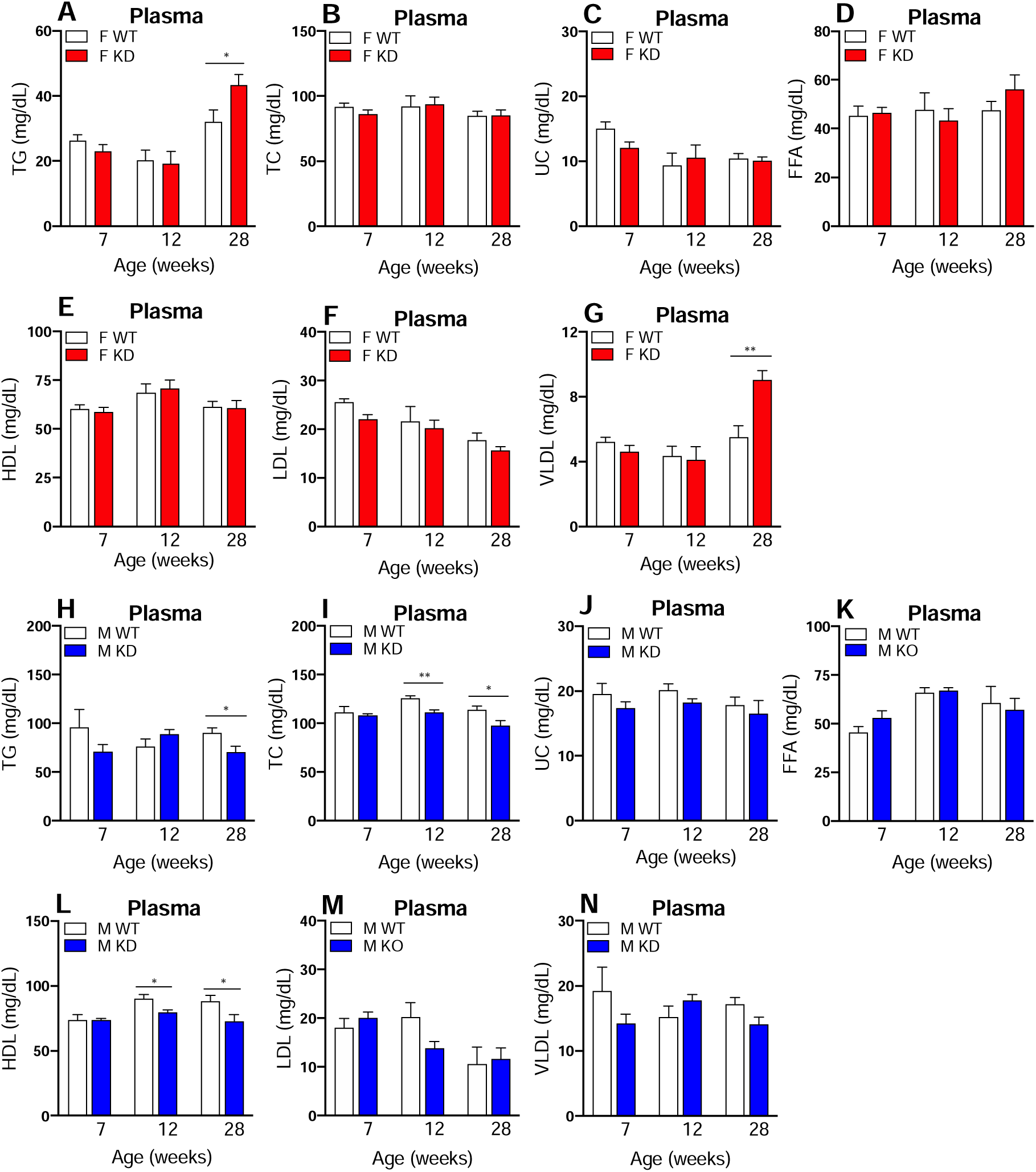
Lipid profiling quantification in *Glo1^+/-^* mice. Plasma lipoproteins **(A)** TG, **(B)** TC, **(C)** UC, **(D)** FFA, **(E)** HDL, **(F)** LDL and **(G)** VLDL were quantified in fasted female mice at 7 (n=9), 12 (n=9) and 28 (n=9) weeks of age. Plasma lipoproteins **(H)** TG, **(I)** TC, **(J)** UC, **(K)** FFA, **(L)** HDL, **(M)** LDL and **(N)** VLDL were quantified in fasted male mice at 7 (n=9), 12 (n=9) and 28 (n=9) weeks of age. Data is presented as mean ± SEM and statistical significance among groups was calculated by Student t-test. *p<0.05, **p<0.01, ***p<0.001.

### 3.5 Reduced *Glo1* levels do not result in atherosclerosis

To assess if *Glo1^+/-^* resulted in cardiovascular phenotypes, we evaluated the occurrence of atherosclerotic lesions in the aortic arch using Red Oil O staining. Our results showed no evidence for aortic lesions in 34-week-old female and male *Glo1^+/-^* mice (n=4-5/group), suggesting that partial loss of *Glo1* alone does not induce atherosclerosis in mice. These findings were not surprising given the natural resistance of atherosclerosis in B6 mice, as our mice were not crossed with atherosclerosis-prone backgrounds such as *ApoE*^-/-^ or *Ldlr*^-/-^, nor were they challenged with atherogenic diets.

### 3.6 *Glo1^+/-^* exhibits perturbed lipid pathways in metabolic tissues

To assess if lipid pathways were altered in parallel with the lipid phenotypes observed in *Glo1^+/-^* mice, we assessed select fatty acid and triglyceride metabolism genes in liver, gonadal adipose and skeletal muscle in mice of both sexes using qPCR (*Lipin1, Acc1, Fasn, Elovl6, Scd1, Srebp1c, Dgat1 and Dgat2)*. We found both sex-specific alterations and expression changes common to both sexes in these genes (Figure 4A). In *Glo1^+/-^* mice, both sexes exhibited decreased expression of *Acc1*, encoding the rate-limiting enzyme in fatty acid synthesis, across all tissues; increased expression of *Fasn*, encoding a key enzyme in *de novo* lipogenesis, in gonadal adipose tissue; and increased expression of *Dgat2*, encoding the enzyme that directly catalyzes the formation of triglycerides, in gonadal adipose tissue. Importantly, the concurrent upregulation of *Fasn* and *Dgat2* in gonadal adipose is consistent with the obesity phenotype observed in *Glo1^+/-^* mice. Genes that exhibited sex-specific alterations included *Fasn*, *Dgat1* and *Scd1* in various metabolic tissues*. Dgat1,* which directly catalyzes the formation of triglycerides, was upregulated in female gonadal adipose (p=0.0394) but not male gonadal adipose (p=0.73) despite the increased gonadal adipose fat pad in male *Glo1^+/-^* mice (Figure 1O; male *Glo1^+/-^*p=0.0314). Male *Glo1^+/-^* mice showed significantly upregulated *Scd1* in liver and skeletal muscle and no difference in gonadal adipose. Scd1 is the rate-limiting enzyme in the biosynthesis of monounsaturated fatty acids and protects against adiposity when knocked out in mice [38]. It is possible that the upregulation of *Scd1* in male *Glo1^+/-^* mice in skeletal muscle may contribute to further adiposity within the tissue. Females *Glo1^+/-^* mice did not show any difference in *Scd1* expression but had significantly increased *Fasn* in skeletal muscle whereas males did not which further supports the obesogenic and impaired glucose tolerance phenotype specific to females [39]. In short, these results not only indicate sex-specific alterations of the lipid metabolic pathways across tissues that may underly sex differences in metabolic phenotypes in *Glo1^+^*^/-^ mice (Figure 4B) but also provide insight into other sex-specific phenotypes observed such as impaired glucose tolerance.

**Figure 4.**
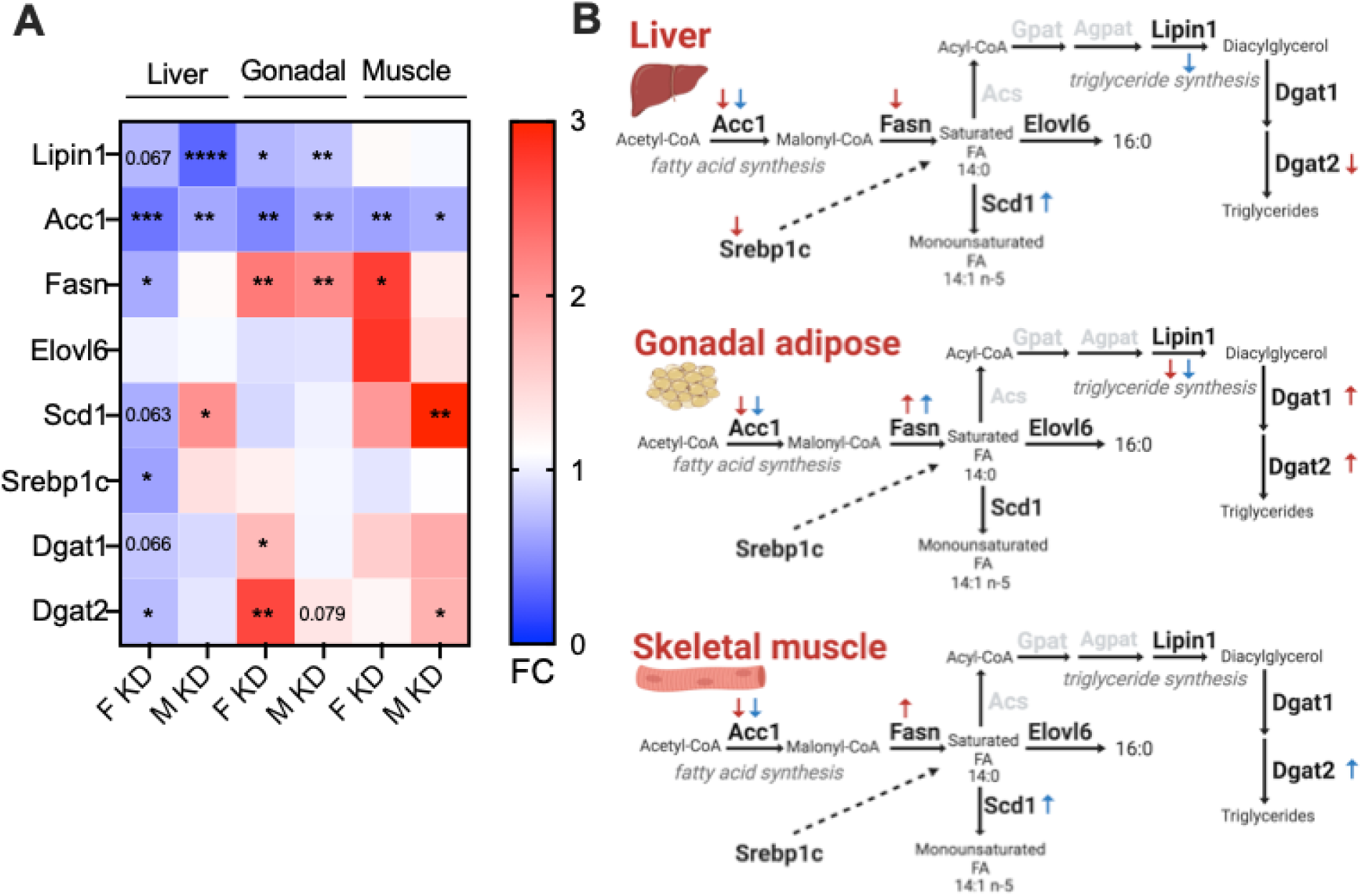
Expression of lipid metabolism genes across liver, gonadal adipose and skeletal muscle tissues in *Glo1^+/-^* mice. **(A)** Expression of lipid metabolism genes (*Lipin1, Acc1, Fasn Elovl6, Scd1, Srebp1c, Dgat1, Dgat2*) were assessed in liver, gonadal adipose and skeletal muscle tissue and in 28-week-old female and male mice (n=4/group). **(B)** Lipid metabolism pathway in liver, gonadal adipose and skeletal muscle tissue is shown to reflect significant data in heatmap with red arrows depicting female Glo1*^+/-^* mice and blue arrows for male Glo1*^+/-^* mice. Arrows pointing up depicts significantly upregulated mRNA expression and arrows pointing down represent significantly downregulated mRNA expression. Genes that encode for enzymes that are part of the lipid metabolism pathway but not tested are represented by the grey color (*Acs, Gpat, Agpat*). Data was analyzed using unpaired t-test with Welch’s correction. Statistically significant data is denoted by *p<0.05, **p<0.01, ***p<0.001, ****p>0.0001.

### 3.7 AGE accumulation does not explain the metabolic changes in *Glo1^+/-^* mice

AGEs have been implicated as a mediator of Glo1-induced metabolic disorders [1, 4, 10]. To explore whether *Glo1^+/-^* phenotypes and lipid pathway reprogramming in metabolic tissues were a result of MG-derived AGEs, MGH1, CEL and other AGE-related components were assessed in metabolic tissues by ELISA, qPCR and microarray (Figure 5A-5E). MGH1 and CEL are both MG-derived AGEs that bind to Rage. MGH1 and CEL are two of the most abundant AGEs, making both compounds a suitable choice for AGE analysis [40–42]. MGH1 (n=6-8/group) and CEL (n=5/group) were quantified in the liver, gonadal adipose and skeletal muscle in both sexes (Figure 5B). In *Glo1^+/-^* females, there was no significant difference in MGH1 and CEL levels in any of the tissues analyzed (Figure 5B). In contrast, MGH1 was significantly elevated in male *Glo1^+/-^* skeletal muscle compared to controls while no differences were observed for CEL (Figure 5B). Overall, our results suggest that the metabolic disturbance induced by Glo1 reduction in mice, particularly in females, may not be solely mediated through changes in AGEs.

**Figure 5.**
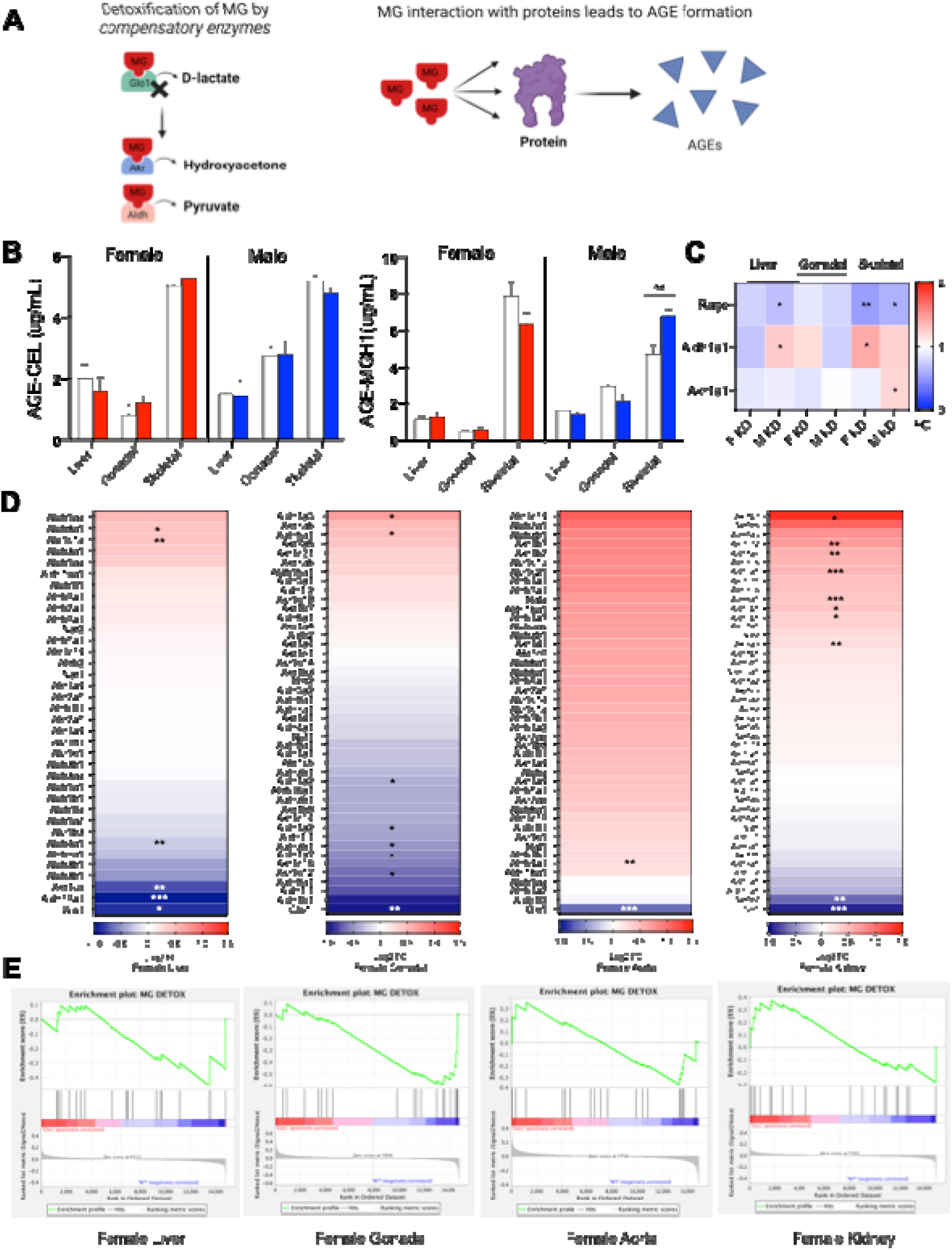
Quantification of AGEs and AGE-related components across liver, gonadal adipose and skeletal muscle tissues in *Glo1^+/-^* mice. **(A)** Schematic overview of MG detoxification mechanism and the generation of AGEs Quantification by ELISA of **(B)** CEL and MGH1 in male and females at 28 weeks of age across metabolic tissues including liver, gonadal adipose and skeletal muscle (n=6-7/group). Gene expression data by qPCR for **(C)** *Ager*, *Akr1a1* and *Aldh1a1* across liver, gonadal adipose and skeletal muscle tissues (n=5-6/group). **(D)** Gene expression data by microarray for Akr and Aldh subtypes in female liver, gonadal adipose, aorta and kidney at 34 weeks of age. (**E)** GSEA plots of gene expression data by microarray for Akr and Aldh subtypes in female liver, gonadal adipose, aorta and kidney at 34 weeks of age. Data was analyzed using unpaired t-test with Welch’s correction. Statistically significant data is denoted by *p<0.05, **p<0.01, ***p<0.001, ****p>0.0001.

To further investigate the role of AGEs in modulating metabolic dysfunction in *Glo1^+/-^* mice, gene expression for *Ager,* the gene that encodes for receptor for advanced glycation endproducts (Rage), was assessed in liver, gonadal adipose and skeletal muscle of mice (Figure 5C). *Ager* was significantly downregulated in both female and male *Glo1^+/-^*mice in liver and skeletal muscle tissues compared to their corresponding controls, while it showed a downregulated trend in *Glo1^+/-^* adipose. The downregulated expression of *Ager* across tissues in both sexes also suggests that the metabolic phenotypes may not be solely modulated by AGEs. To further explore the potential role of Rage signaling in mediating the metabolic phenotypes observed, downstream gene expression of Rage signaling was assessed in liver, gonadal adipose tissue and kidney using microarray (Supplementary Figure 3A-3D). These findings revealed that the genes involved in Rage signaling such as ROS signaling genes, Nfkb signaling genes and apoptosis signaling genes were not consistently and significantly altered between *Glo1^+/-^* mice and wildtype groups in the tissues assessed. This further supports the unlikely role of Rage signaling activation or AGEs in mediating the metabolic alterations observed in *Glo1^+/-^* mice.

Due to the underwhelming evidence AGE involvement or Rage signaling observed in female and male *Glo1^+/-^* mice, we suspected that Glo1 compensatory enzymes such as Ark and Aldh [17] were likely involved in metabolizing MG, thus preventing the formation of AGEs and subsequent Rage signaling activation. To investigate the role of these compensatory enzymes in Glo1*^+/-^* mice, we assessed gene expression of *Akr1a1* and *Aldh1a1* in liver, gonadal adipose and skeletal muscle tissues (Figure 5C). *Ark1a1* was significantly upregulated in male *Glo1^+/-^* skeletal muscle without any changes in the other tissues. *Aldh1a1* was also significantly upregulated in male *Glo1^+/-^*liver and female *Glo1^+/-^* skeletal muscle. Neither of the compensatory enzymes were altered in gonadal adipose tissue. To supplement these results, we looked at the gene expression changes of the numerous *Akr* and *Aldh* subtypes in female liver gonadal adipose, aorta and kidney (Figure 5D, 5E). Surprisingly, these results revealed that several *Akr* and *Aldh* subtypes were significantly upregulated, particularly in the Glo1*^+/-^* female kidney compared to the female controls *(Akr1b3, Akr1c12, Aldh1a7, Aldh8a1, Aldh3b1* and *Akr7a5*). Increased compensatory enzymes in the kidney may suggested heightened detoxification of MG taking place in the kidney.

Together these results, which include the lack of changes in AGEs levels in all female tissues and most male tissues, the downregulation of *Ager* (which is key to AGE/RAGE signaling pathway) across most tissues in both sexes, and the upregulation of Glo1 compensatory enzymes (that prevent AGE formation) in select tissues, do not support a major role of AGEs in the observed metabolic effects in Glo1*^+/-^*. Importantly, the upregulation of Glo1 compensatory enzymes, particularly in the female kidney, can detoxify MG to metabolites such as pyruvate, a key metabolite that can regulate metabolism.

### 3.8 *Glo1^+/-^* exhibit tissue-specific transcriptomic alterations in multiple metabolic tissues

To uncover the molecular basis of the obesity and glucose intolerance in female *Glo1^+/-^* mice, we profiled the transcriptome of metabolic tissues including gonadal adipose, aorta, and liver in 34-week-old female mice. At FDR < 0.1, a total of 93, 347, 421, 232 differentially expressed genes (DEGs) were found in adipose tissue, aorta, liver, and kidney respectively (Figure 6A; Table 1), suggesting large-scale transcriptomic alterations in *Glo1^+/-^* mice.

**Figure 6.**
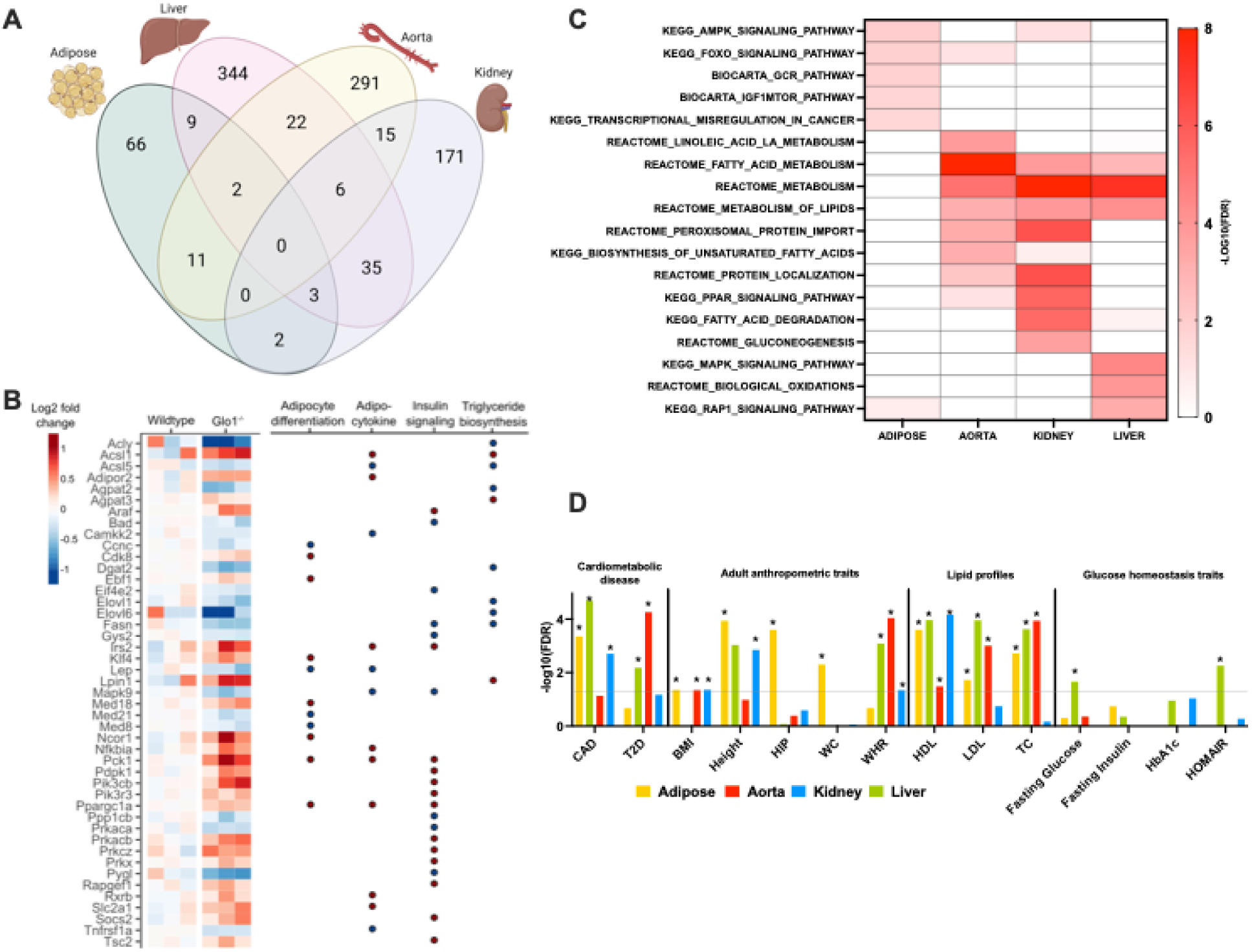
Summary of differentially expressed gene signatures in *Glo1^+/-^* mice. **(A)** Venn diagram of the number of significant DEGs with FDR < 10% in adipose, aorta and liver. **(B)** Individual sample expression level changes for genes involved in adipocyte differentiation, adipocytokine and insulin signaling with significant expression level difference with P < 0.05. Fold change was determined by comparing the expression values of Glo1^+/-^ mice to the mean value of wildtype samples. Red and blue indicate increased and decreased expression in knockdown mice, respectively. **(C)** Canonical pathways with enrichment FDR < 5% are shown. Statistical significance was determined by Fisher’s exact test. **(D)** *Glo1* reduction in female mice and genetic association to human metabolic diseases. For human metabolic trait association, statistical significance was determined by MSEA. Asterisks indicate a significant association with FDR<5%. CAD, coronary artery disease; T2D, type 2 diabetes; BMI, body mass index; HIP, hip circumference; WC, waist circumference; WHR, waist-hip ratio.

**Table 1.** Top 15 transcription factors whose downstream targets are enriched for Glo1*^+/^*DEG signatures in adipose tissue, aorta, liver and kidney.

| Adipose |  | Aorta |  | Liver |  | Kidney |  |
| --- | --- | --- | --- | --- | --- | --- | --- |
| TF | P | TF | P | TF | P | TF | P |
| **NR3C1** | 1.97E-06 | **PPARG** | 1.12E-07 | **POLR2B** | 1.34E-04 | **NR1D1** | 9.55E-06 |
| **PPARG** | 3.03E-06 | **NR3C1** | 8.30E-06 | **H2AZ** | 1.49E-04 | **POLR2B** | 6.64E-05 |
| **HDAC3** | 1.97E-05 | **PPARA** | 1.24E-05 | **XBP1** | 2.04E-04 | **HNF4A** | 7.13E-05 |
| **PPARA** | 3.52E-05 | **HDAC3** | 2.19E-05 | **NR1D1** | 2.14E-04 | **RXRA** | 9.92E-05 |
| **RXRA** | 4.54E-05 | **RXRA** | 9.31E-05 | **KDM6A** | 3.21E-04 | **XBP1** | 1.19E-04 |
| **CEBPB** | 1.41E-04 | **HNF4A** | 9.92E-05 | **RORA** | 4.09E-04 | **RORA** | 1.84E-04 |
| **EP300** | 1.84E-04 | **NR1D1** | 1.66E-04 | **KMT2C** | 4.59E-04 | **ARNTL** | 1.84E-04 |
| **CEBPA** | 4.26E-04 | **PRDM16** | 2.14E-04 | **MYC** | 5.51E-04 | **NR1H2** | 2.35E-04 |
| **STAT5B** | 6.11E-04 | **CRY2** | 2.24E-04 | **CREB1** | 6.11E-04 | **CRY2** | 2.70E-04 |
| **NR1H2** | 6.11E-04 | **NR1H2** | 2.82E-04 | **HDAC3** | 6.32E-04 | **ARID1A** | 3.35E-04 |
| **HNF4A** | 8.97E-04 | **PER2** | 3.08E-04 | **CLOCK** | 6.98E-04 | **PPARA** | 4.42E-04 |
| **STAT5A** | 8.97E-04 | **ESR2** | 3.08E-04 | **PHF8** | 7.21E-04 | **HDAC3** | 4.77E-04 |
| **NCOR1** | 9.52E-04 | **CEBPB** | 3.63E-04 | **GTF2B** | 8.70E-04 | **MYC** | 4.77E-04 |
| **ESR1** | 1.16E-03 | **CRY1** | 3.63E-04 | **CRY2** | 1.01E-03 | **CLOCK** | 4.94E-04 |
| **RXRG** | 1.26E-03 | **CEBPA** | 5.13E-04 | **RXRA** | 1.19E-03 | **KDM6A** | 5.70E-04 |

*Glo1^+/-^* gene signatures exhibited strong tissue specificity when we compared the DEGs across tissues (Figure 6A). For adipose tissue, aorta, liver, and liver, 71.0%, 83.9%, 81.7%, and 73.7% of signatures were unique for each respective tissue. Two genes were consistently down-regulated in adipose, liver, and aorta, including *Nqo2* (N-Ribosyldihydronicotinamide: Quinone Reductase) and *Tuba1a* (Tubulin Alpha 1a). *Nqo2* is an antioxidant that may protect against diabetes-induced endothelial dysfunction [43], whereas *Tuba1a* is the gene responsible for producing cytoskeleton protein tubulin, a known target for glyoxalase system defect [5]. The consistent decreases in these genes suggest that reduced *Glo1* levels promote oxidative stress and damage to cellular structure.

### 3.9 *Glo1* reduction perturbs metabolic pathways systemically

To determine the pathways affected by *Glo1* reduction in each tissue, we performed pathway enrichment analysis of the tissue-specific DEGs from female *Glo1^+/-^* mice (Figure 6B). We found an enrichment of genes involved in insulin resistance, insulin-like growth factor binding protein complex, monocarboxylic acid metabolic process, which is consistent with the glucose intolerance phenotype in *Glo1^+/-^* female mice [44–47]. Additionally, dysregulation in pathways related to lipid homeostasis were consistently captured across adipose (AMPK signaling, IGF1MTOR pathway, GCR pathway), aorta (fatty acid metabolism, metabolism of lipids, biosynthesis of unsaturated fatty acids), kidney (PPAR signaling, fatty acid degradation), and liver (MAPK signaling, biological oxidations). These results suggest that *Glo1* reduction has a systemic impact on metabolic pathways across tissues. Furthermore, we specifically investigated the fold change of expression of the DEGs involved in adipocyte differentiation, adipocytokine, insulin signaling and fatty acid/triglyceride biosynthesis in adipose tissue (Figure 6C). We observed alterations in key genes involved these processes such as *Elovl6, Fasn, Lep, Ppargc1a, Mapk9*, and *Pck1*. These transcriptomic changes in female mice may underlie the obesity, hyperlipidemia, and glucose intolerance phenotypes observed in females *Glo1^+/-^* mice.

### 3.10 *Glo1* reduction affects genes that show genetic association to human metabolic diseases

Although *Glo1* has been previously identified as a candidate gene for human metabolic disorders [1, 8–10], the underlying mechanisms have not been explored. To assess our previous findings (Figure 6A-6C) and its potential link to human health, we took advantage of the publicly available summary statistics from large-scale human GWAS and used the Marker Set Enrichment Analysis (MSEA) from the Mergeomics package [37] to evaluate whether the *Glo1* signature genes in individual tissues exhibited an over-representation of human disease risk variants [22]. As shown (Figure 6D), the strongest and most consistent associations of suggestive DEGs across tissues were found for three lipid traits (LDL, HDL, and TC). We also identified a significant association of the Glo1 DEGs with BMI, height and waist/hip ratio (WHR). Additionally, consistent with hyperglycemia and dampened glucose tolerance *Glo1^+/-^*, the liver-specific signatures are associated with fasting glucose, HbA1c and HOMA-IR in human GWAS. Moreover, we observed strong association of *Glo1* DEGs in different tissues with CAD and T2D. These results indicate that *Glo1* reduction perturbs genes that have been associated with various human metabolic phenotypes or diseases.

### 3.11 Identification of transcription factor (TF) hotspots underlying *Glo1^+/-^* signatures

To better understand how *Glo1* reduction leads to the perturbations of DEGs in individual tissues, we explored transcription factors (TF) that may mediate the gene expression changes induced by *Glo1* knockdown. We utilized the computational tool BART to predict TF enrichment for Glo1*^+/-^* signatures. Interestingly, PPARg, the master regulator of adipogenesis, is among the top perturbed adipose TF hotspots, as its target genes are highly enriched among the adipose DEGs (Table 1). Moreover, several other TFs implicated directly or indirectly in adipogenesis were also highly ranked in adipose TF hotspots, including CEBPB/A, PPARA, HNF4A, and RXRA. These findings align with the increased adiposity in female *Glo1^+/-^*mice. In aorta, we identified perturbations in a number of TFs involved in circadian clock regulation (CRY1, CRY2, PER2) [48], vascular inflammation (PPARG, NR3C1) [49], and those linked in coronary artery disease, including the estrogen receptor, ESR2. In the liver, several TFs are related to circadian clock regulation (CLOCK, CRY2, NR1D1), liver metabolism (NR1D1, RORA) and liver development (KDM6A, MYC, HDAC3), indicating the potential of Glo1 to alter a wide spectrum of physiological activities in the liver. Lastly, in the kidney following a similar trend to the liver we highlight various circadian TFs (CLOCK, CRY2, ARNTL, NR1D1), kidney development TFs (POLR2B, HNF4A, HDAC3, MYC, KDM6A) and those involved in glucose and lipid metabolism (NR1H2, PPARA, RORA). With sex differences being present in our phenotypic results, we looked to highlight at the TF level if there are any known TFs that have a female bias to potentially explain these differences. Of interest, when comparing our results to known sex biased TFs from the Gene by Tissue Expression (GTEx) study [50], we highlight that across each tissue that a minimum of 22% of our TFs have evidence of being female biased: adipose (16/60, 27%), aorta (14/64, 22%), liver (11/44, 25%) and kidney (9/38, 24%). Importantly, a number of these female biased TFs are related to circadian rhythm (ARNTL, NR1D1), tissue development (HNF4A) and metabolism (PPARA, HNF4A), which collectively may in part explain female-specific phenotypic differences including poorer glucose tolerance. These tissue-specific TFs likely mediate the effects of *Glo1* reduction on the molecular pathways altered in individual tissues.

### 3.12. Inference of the potential role of MG metabolite pyruvate in regulating *Glo1^+/-^* signatures

To further explore regulatory molecules that reprogram the transcriptome, we focused on MG metabolites produced by compensatory enzymes that we found to be upregulated in our gene expression analysis. In particular, pyruvate can be produced from MG detoxification which can further regulate gene expression. Indeed, we found that the genes involved in gluconeogenesis (*Pcx, Pck, Enol, Pgam1, Pgk1, Gapdh, Fbp1* and *G6pc*) were mostly upregulated in female *Glo1^+/-^* kidney tissues and to a lesser extent in female *Glo1^+/-^* liver, adipose and aorta tissues (Figure 7). These results suggest a potential role of MG metabolites in regulating gene expression and glucose production.

**Figure 7.**
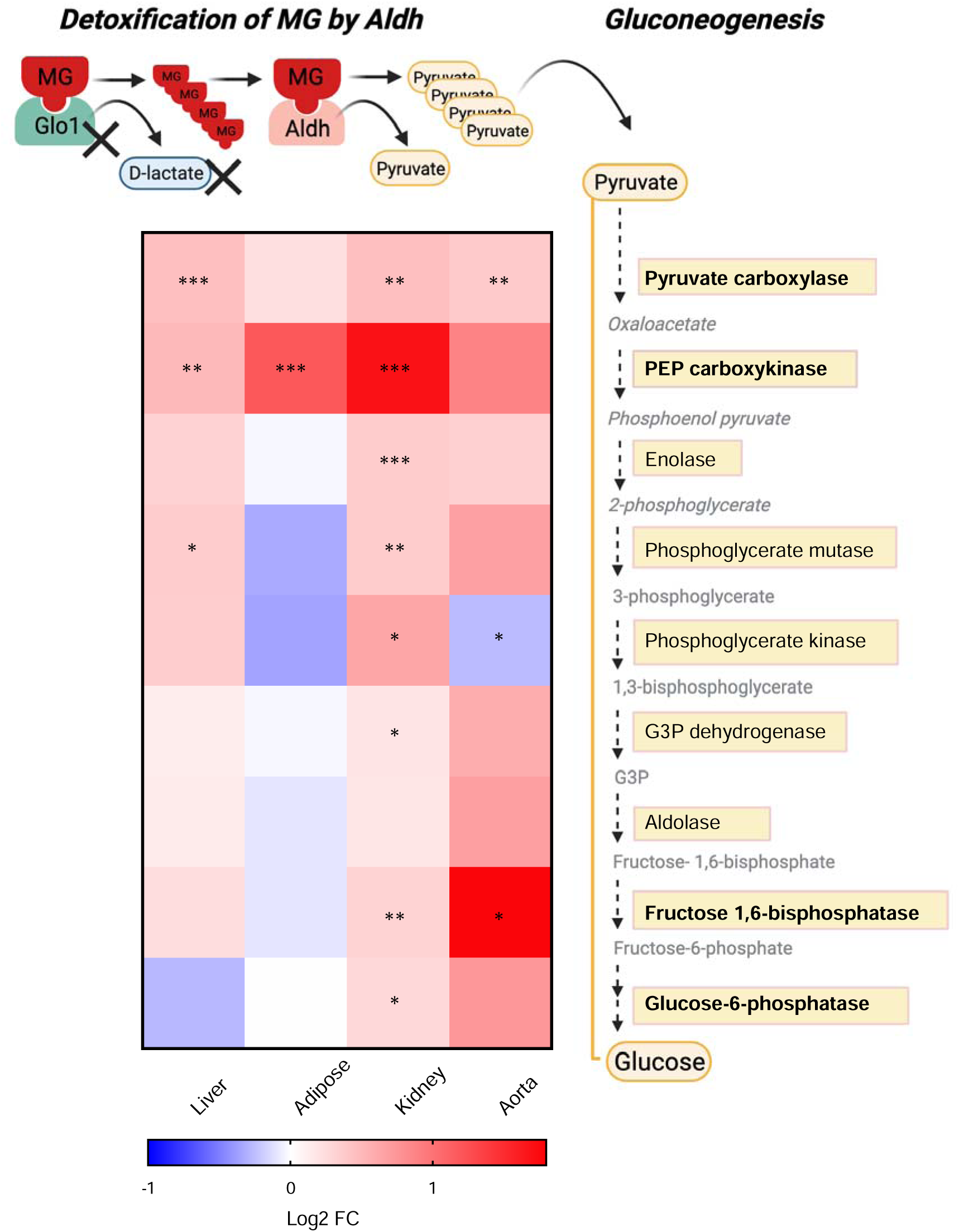
Schematic summary of gluconeogenesis pathway showing the potential role of MG metabolite pyruvate in regulating *Glo1^+/-^* signatures. Schematic overview of MG detoxification by Aldh in the kidney leading to the generation of pyruvate. Metabolic intermediates such as pyruvate can enter the gluconeogenesis pathway resulting in the conversion of pyruvate to glucose. Asterisks indicate a significant increase in gene expression of enzymes involved in gluconeogenesis.

## 4. DISCUSSION

To explore the role of *Glo1* in metabolic disorders, we conducted a systems biology study of *Glo1^+/-^* mice. We carried out extensive metabolic phenotyping in both sexes at various ages and further performed transcriptomic and AGE profiling of key metabolic tissues. This was followed by network and integrative genomics analyses to elucidate the underlying regulatory cascades and human disease relevance (Figure 8). From this study, we found that *Glo1^+/-^* mice demonstrated metabolic dysregulation, including increased body weight and fat mass in both sexes in adulthood. Metabolic dysregulation was also observed in a sex-dependent manner, including increased muscle mass, hypoglycemia as well as decreased TC, TG and HDL in male *Glo1^+/-^* mice, whereas increased circulating TG and VLDL and impaired glucose tolerance was observed in female *Glo1^+/-^*mice. The metabolic phenotypes observed in *Glo1^+/-^* female mice were paralleled by large-scale tissue-specific expression alterations in metabolic genes and pathways across tissues. Select lipid metabolic pathway alterations were also confirmed by qPCR experiments in both males and females. The tissue-specific molecular signatures were further tied to numerous tissue-specific TFs that regulate adipogenesis, insulin signaling, aortic endothelial functions, and key liver functions, and collectively demonstrated significant association with multiple human metabolic traits. To explain the source for the observed systemic effects of *Glo1* reduction, we quantified the most prominent AGEs (CEL and MGH1) in metabolic tissues and found AGE levels did not differ between *Glo1^+/-^* mice and WT controls in most tissues. This led our investigation to assess potential changes in AGE receptor gene expression, namely *Ager*, finding no significant differences between *Glo1^+/-^*mice and WT mice (Figure 5C). Our findings largely agree with those from the Wortmann et al. study which did not detect a difference in AGEs in heart, kidney, and liver between *Glo1^+/-^* and WT [55]. Overall, the phenotypes observed in our specific *Glo1^+/-^*mice cannot be explained by the AGE pathway as a major factor in our study.

**Figure 8.**
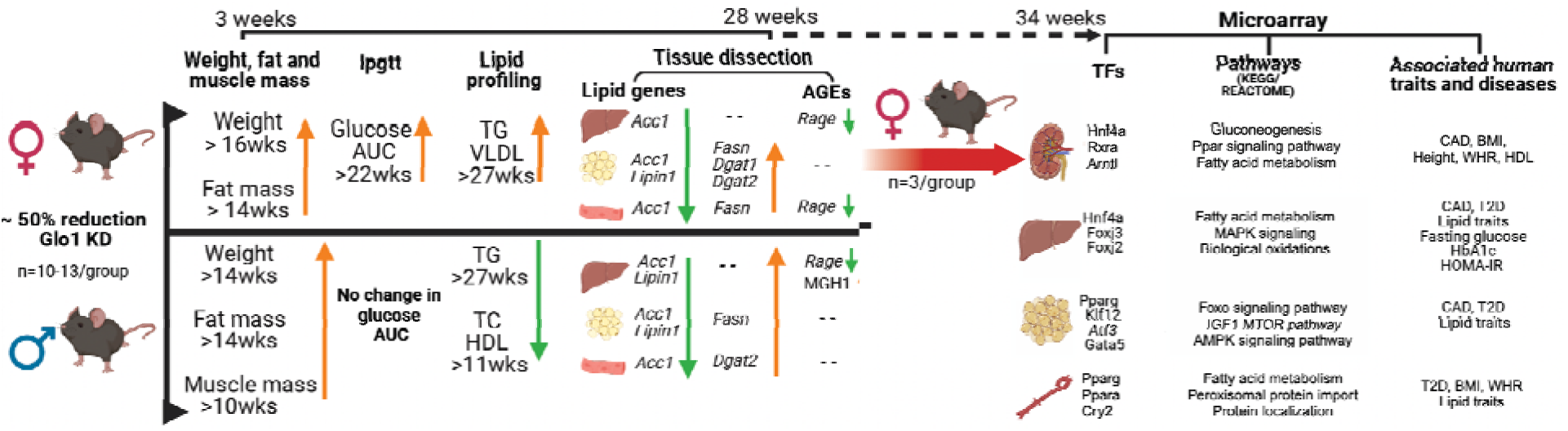
Schematic summary of *Glo1* deficit effects in female and male mice. Schematic overview of study design showing the metabolic phenotypes induced by *Glo 1* reduction in female and male mice over time.

Our observation that *Glo1^+/-^* exhibited increased weight and adiposity is consistent with the previous linkage study that revealed *Glo1* resided under a QTL for body weight in mice [51]. In females, the body weight increases in *Glo1^+/-^*mice was largely due to growth of various adipose depots since lean mass remained on par with control mice, whereas increased body weight in male *Glo1^+/-^*mice were attributed to both fat and muscle mass increase. Lipid metabolism genes *Fasn* and *Dgat2* were upregulated in female and male *Glo1^+/-^*mice in gonadal adipose, supporting the obesogenic phenotypes observed (Figure 1, Figure 4). Along with the obesity phenotypes, female *Glo1^+/-^* mice also demonstrated comorbidities such as glucose intolerance. As shown in our data (Figure 7), generation of metabolites such as pyruvate that arise from MG clearance by compensatory enzymes such as Aldh may promote reprogramming of transcriptome may also contribute to the dysregulated glucose phenotypes in female *Glo1^+/-^*mice. Surprisingly, male *Glo1^+/-^* mice showed decreased plasma glucose. A possible explanation for this may be due to increased uptake of circulating glucose by the increased skeletal muscle tissue in male *Glo1^+/-^*mice (Figure 1). These unique phenotypes demonstrated by male *Glo1^+/-^* mice may explain the enhanced ability of male *Glo1^+/-^* mice to adequately regulate blood glucose levels by removing excess glucose from the circulation. Further assessment of changes in glucose metabolism in male *Glo1^+/-^* skeletal muscle is warranted. In addition, female *Glo1^+/-^* mice had significantly increased *Fasn* in skeletal muscle whereas males did not, which further supports the obesogenic and impaired glucose tolerance phenotype in female but not male *Glo1^+/-^* mice. Lastly, along with obesity, the female *Glo1^+/-^* mice also demonstrated additional comorbidities such as hyperlipidemia with increased TG and VLDL compared to WT mice. Male *Glo1^+/-^* mice also showed a dysregulated lipid profile with significantly decreased TG, TC, and HDL compared to WT mice.

Interestingly, many of the perturbed metabolic profiles were observed after 12 weeks of age, suggesting an age-dependent effect in *Glo1^+/-^*mice. This observation is consistent with the lack of increased body weight previously reported for the *Glo1^+/-^* by Wortmann et al. using the same mouse model where the follow-up period was only 10 weeks [52]. This implicates the potential of partial deficiency in programming the body towards a state prone to metabolic dysfunctions in late adulthood, which heightens the risk of age-associated metabolic diseases.

Investigation of the female adipose transcriptome revealed changes in many pathways and key regulatory genes involved in adipocyte differentiation, adipocytokine signaling, insulin signaling, fatty acid/triglyceride biosynthesis, and PPAR signaling, a state that promotes adipogenesis and subsequent obesity risk [53, 54]. This is further supported by the identification of adipogenic master regulators including PPARG as TF hotspots for the DEGs altered in *Glo1^+/-^* female mice. The liver gene signatures were significantly enriched for fatty acid metabolism along with bile acid metabolism and glucose metabolism, consistent with the altered TG, VLDL levels and insulin sensitivity in *Glo1^+/-^* female mice. As males exhibited decreased lipid profiles along with lower fasting glucose levels, we tested genes involved in lipid metabolism in both sexes using qPCR and found disparate tissue-specific changes in numerous genes (*Fasn*, *Scd1*, *Srebp1c*, etc.) between sexes, providing molecular support for the distinct phenotypic manifestations between males and females. However, we did observe consistent changes in genes such as *Acc1* (fatty acid synthesis) and *Lipin1* (triglyceride synthesis). The different lipid profiles between sexes likely reflect the collective differential activities of the lipid metabolism pathways across tissues between males and females (Figure 4A).

Our characterization of sex-specific responses in metabolic traits yielded novel results suggesting that female mice are more vulnerable to *Glo1* reduction, especially in maintaining glucose and lipid homeostasis. We also revealed that 4 out of 17 genes (*Ebp*, *Lss*, *Hsd17b7*, *Nsdhl*) in the steroid synthesis pathway have altered expression exclusively in female adipose tissue (Fold change = 6.3, p = 2.6e-4). As a critical pathway that modulates sexual dimorphic adiposity, disruption of steroid synthesis may serve as a mediator for the effect of reduced Glo1 on adiposity in females. However, we only examined the full transcriptome in females and only tested select lipid genes in both sexes in the current study, therefore, further investigations are warranted to assess additional sexually dimorphic pathways between sexes. However, some explanation to the female specific vulnerability to glucose intolerance can be explained through our uncovering of known female bias TFs. These TFs are critical to circadian rhythms, which in turn plays a role in glucose metabolism as well as those TFs such as HNF4A, which have dual roles in development and gluconeogenesis. A likely increase in gluconeogenesis may be feasible highlighted by increases in numerous genes such as *G6pc* and *Pck* involved in this process, particularly in female tissues (Figure 7) as well as a significant enrichment in gluconeogenesis in the kidney (Figure 6C).

*Glo1* has been previously implicated as a key regulator for CAD [8]. We did not observe direct evidence for atherosclerotic lesions in 34-week-old *Glo1*^+/-^ mice, likely due to the natural resistance of mice for atherogenesis without extreme genetic and dietary perturbations. However, the transcriptomic profiling of aorta provided supportive links between *Glo1* and atherosclerosis. Pathway analysis of the aorta DEGs showed significant alterations related to lysosome and lipid metabolism, processes implicated in atherogenesis [55, 56]. Additionally, TF prediction for the aorta DEGs identified TFs that are related to circadian rhythms and endothelial inflammation, which are also important processes for atherogenesis [49], [57]. Furthermore, the Glo1 DEG signatures from multiple tissues examined were also strongly associated with lipid profiles (TC, LDL, and HDL) and CAD in human GWAS. The negative atherosclerotic phenotype in *Glo1^+/-^* mice could be due to insufficient reduction of *Glo1* activity, natural resistance to the development of atherosclerotic lesions and no enforced genetic or environmental stress in our study. Previous studies on the role of Glo1 in atherosclerosis have also yielded inconsistent results. Geoffrion et al. showed that *Glo1* transgene or *Glo1^+/-^* did not affect the progression of atherosclerosis in the *ApoE^-/-^* mice at 22 weeks of age, even under diabetic conditions [58]. On the other hand, Jo-Watanabe et al. showed that *Glo1* transgenic mice on *Apoe^-/-^* background under a high-fat diet went through a reduction of glycation and oxidative stress while preventing age-related endothelial dysfunction by the prevention of eNOS inactivation [59]. Future studies that focus on the long-term effects in atherosclerosis susceptible models with significant genetic or environmental challenges could yield more power in elucidating the potential causal role of *Glo1* in atherosclerosis.

These finding suggest that in the presence of a compromised glyoxalase system such as in our *Glo1^+/-^* mice, compensatory mechanisms and enzymes such as Akr and Aldh increase and may reduce the accumulation and effects of MG in the body and explain the lack of prevalent changes in AGEs. While initially, this wards off any immediate effects and consequences of accumulated MG, long-term effects resulting from the alternative methods of detoxifying MG may lead to increased metabolites such as pyruvate which may further reprogram metabolism.

To our knowledge, we are the first to conduct a comprehensive study to show evidence for both sex- and age-dependent effects that result from *Glo1* reduction in mice. In addition, our results highlight numerous TFs that have a known female bias, which may in part explain that female *Glo1^+/-^*mice are more affected than male *Glo1^+/-^* mice, particularly with these biases being found in key pathways such as circadian rhythms and lipid metabolism. The extensive perturbation of numerous metabolic pathways in individual tissues through regulatory genes such as PPARg and HNF4A, which are among the top TFs predicted to regulate the altered genes in *Glo1^+/-^*, implicate *Glo1* in the modulation of these metabolic regulators to affect metabolic risks. Importantly, our integrative genomics analyses coupling the *Glo1* signatures with human genetic studies help connect *Glo1* with numerous human metabolic traits/diseases. These results are largely consistent with the phenotypic profiles observed in mice and substantiate the relevance of *Glo1* to human metabolic diseases.

In summary, our findings support the role of *Glo1* reduction in modulating obesity and metabolic dysfunction in a sex specific manner without strong evidence for the involvement of AGEs. Our discoveries regarding tissue-specific genes, pathways and transcription factor hotspots involved in metabolic regulation downstream of *Glo1* in multiple metabolic tissues support broader molecular functions of Glo1 beyond its known role in the glyoxalase system. Moreover, the application of these findings to human disease further supports the importance of exploring the physiological functions and pathogenic potential of *Glo1*.

## ACKNOWLEDGMENTS

The authors would like to thank Hyaeran Byun, Hannah Qi, Nam Che, Clara Yuhhtman, Sharda Charugundia and Shraddha Rege for technical assistance.

## AUTHOR CONTRIBUTION

X.Y., I.C., and L.S. conceived and designed research; I.C., L.S., I.A., G.D., G.Z., H.Q. and J.S.L. performed experiments; I.C., M.B., L.S., J.S.L., Z.S., R.L., and H.Q. analyzed data, I.C., M.B., L.S., J.S.L. and H.Q. interpreted results of experiments; I.C., M.B., L.S. and J.S.L. prepared figures; X.Y., I.C., M.B., and L.S., drafted manuscript; X.Y., I.C., M.B., L.S., S.W., R.L., and J.S.L. edited and revised manuscript; All authors approved the final version of the manuscript.

