## Supplemental Table 1 and Figures for "Glo1 reduction in mice results in age- and sex-dependent metabolic dysfunction": Glo1 _Supplementary List.docx

**Supplementary Materials List**

**Supplementary Table 1**

qPCR primer list

**Supplementary Figure 1**

Figure 1A: PCR genotype of WT and Glo1 mice at 3 weeks of age

Figure 1B: Glo1 enzyme activity at 28 weeks

**Supplementary Figure 2**

Figure 2A-2D: Food and water intake by WT and Glo1 mice

**Supplementary Figure 3**

Figure 3A: Schematic overview of Rage signaling

Figure 3B: Liver gene expression of Rage signaling

Figure 3C: Gonadal adipose gene expression of Rage signaling

Figure 3D: Kidney gene expression of rage signaling
