## Supplemental Table 1 and Figures for "Glo1 reduction in mice results in age- and sex-dependent metabolic dysfunction": Glo1_supplementary IC1.docx

**Glo1 Supplementary Materials**

**Supplementary Table 1**

List of qPCR mouse primers

| **Gene** | **Forward primers** | **Reverse primers** |
| --- | --- | --- |
| *Glo1* | 5’-GCTTCTCCCACAAGTCTGTG-3’ | 5’-GGTACAGTGCAGGGGAAAGA-3’ |
| *Gapdh* | 5’-AACTTTGGCATTGTGGAAGG-3’ | 5’-ACACATTGGGGGTAGGAACA-3’ |
| *Rage* | 5′-AACACAGGAAGAACTGAAGCTTGG-3' | 5′-CTTTGCCATCGGGAATCAGAAGTT-3′ |
| *Akr1a1* | 5'-GGTATATTGTGCCCATGATTACG-3' | 5'-GGGGAGTAGCAGGCAATG-3' |
| *Aldh1a1* | 5'-GACAGGCTTTCCAGATTGGCTC-3' | 5'-AAGACTTTCCCACCATTGAGTGC-3' |
| *Lipin1* | 5'-CCCTCGATTTCAACGTACCC-3' | 5'-GCAGCCTGTGGCAATTCA-3' |
| *Acc1* | 5'-GGATATCGCATCACAATTGGC-3′ | 5′-CCTCGGAGTGCCGTGCTCTGGATC-3′ |
| *Fasn* | 5'-AGCGGCCATTTCCATTGCCC-3' | 5'-CCATGCCCAGAGGGTGGTTG -3' |
| *Elovl6* | 5’-CCCGAACTAGGTGACACGAT-3’ | 5’-TACTCAGCCTTCGTGGCTTT-3’ |
| *Scd1* | 5'-TTCTTGCGATACACTCTGGTGC-3' | 5'-CGGGATTGAATGTTCTTGTCGT-3' |
| *Srebp1c* | 5'-GGAGCCATGGATTGCACATT-3' | 5'-GGCCCGGGAAGTCACTGT-3' |
| *Dgat1* | 5'-GGAATATCCCCGTGCACAA-3' | 5'-CATTTGCTGCTGCCATGTC-3' |
| *Dgat2* | 5'-CCGCAAAGGCTTTGTGAA-3' | 5'-GGAATAAGTGGGAACCAGATCAG-3' |

**Supplementary Figure 1**

A.

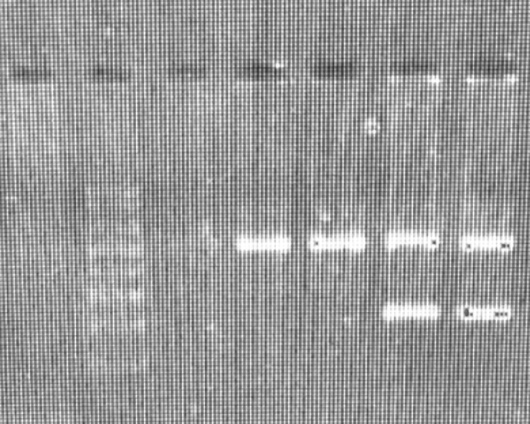

Gapdh (top band)

WT (no band)

Glo 1 KD (bottom band; 220bp)

B.

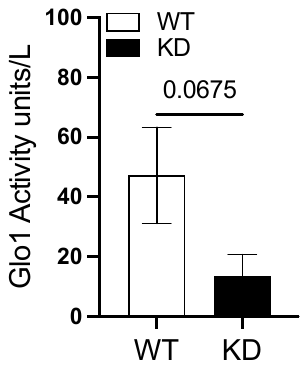

**Supplementary Figure 2**

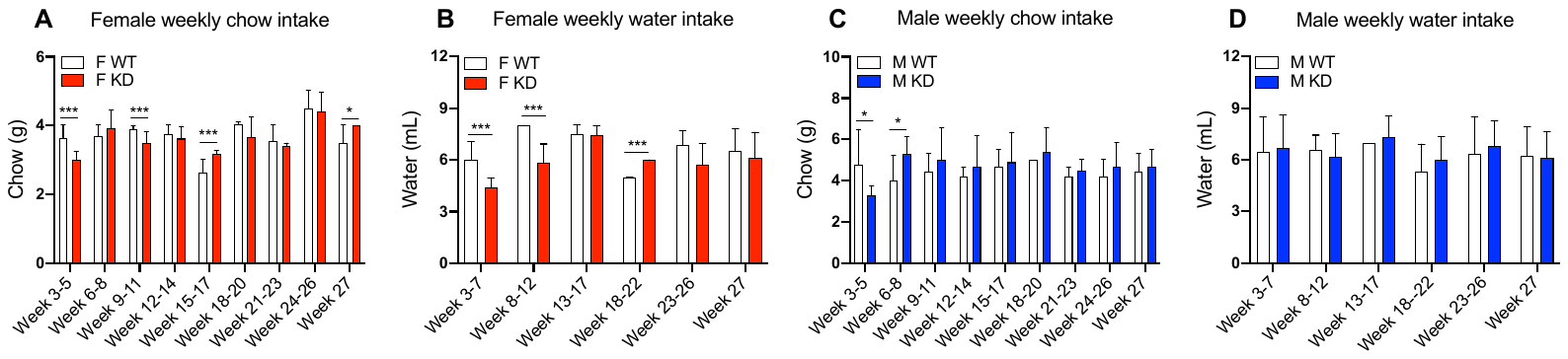

**Supplementary Figure 3**

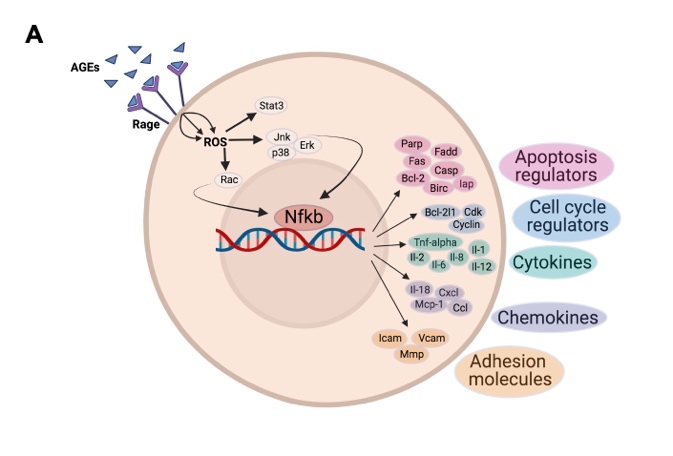

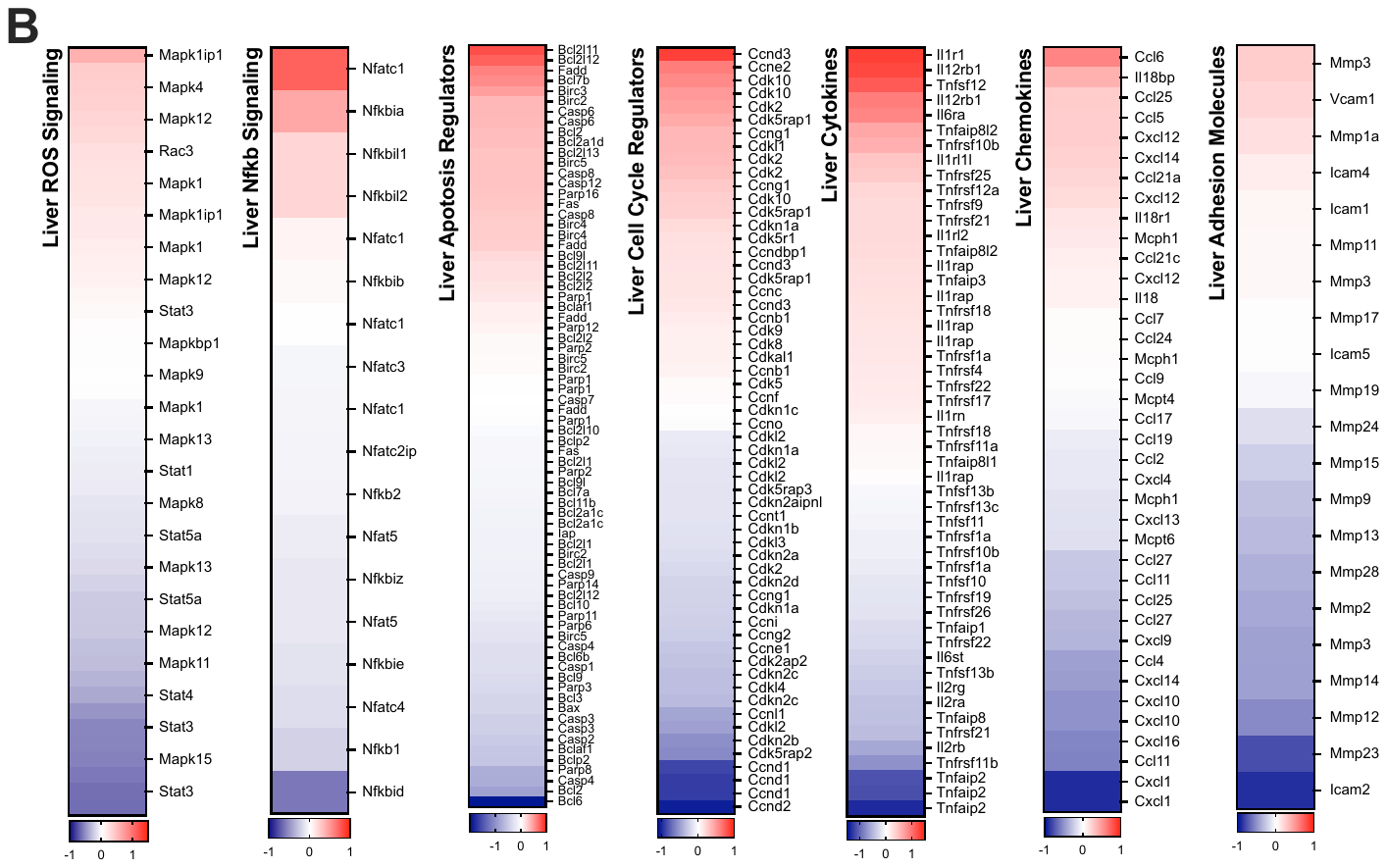

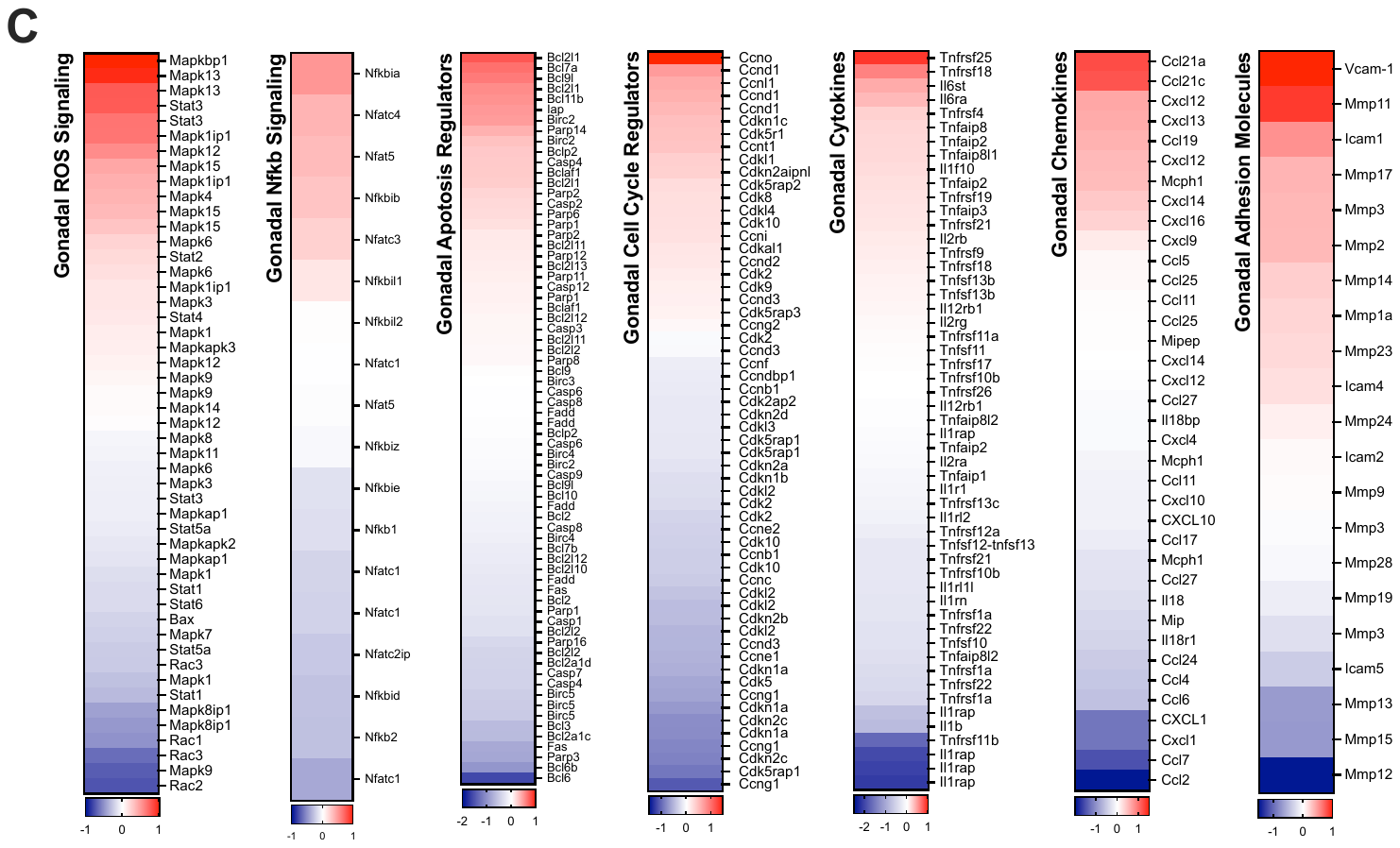

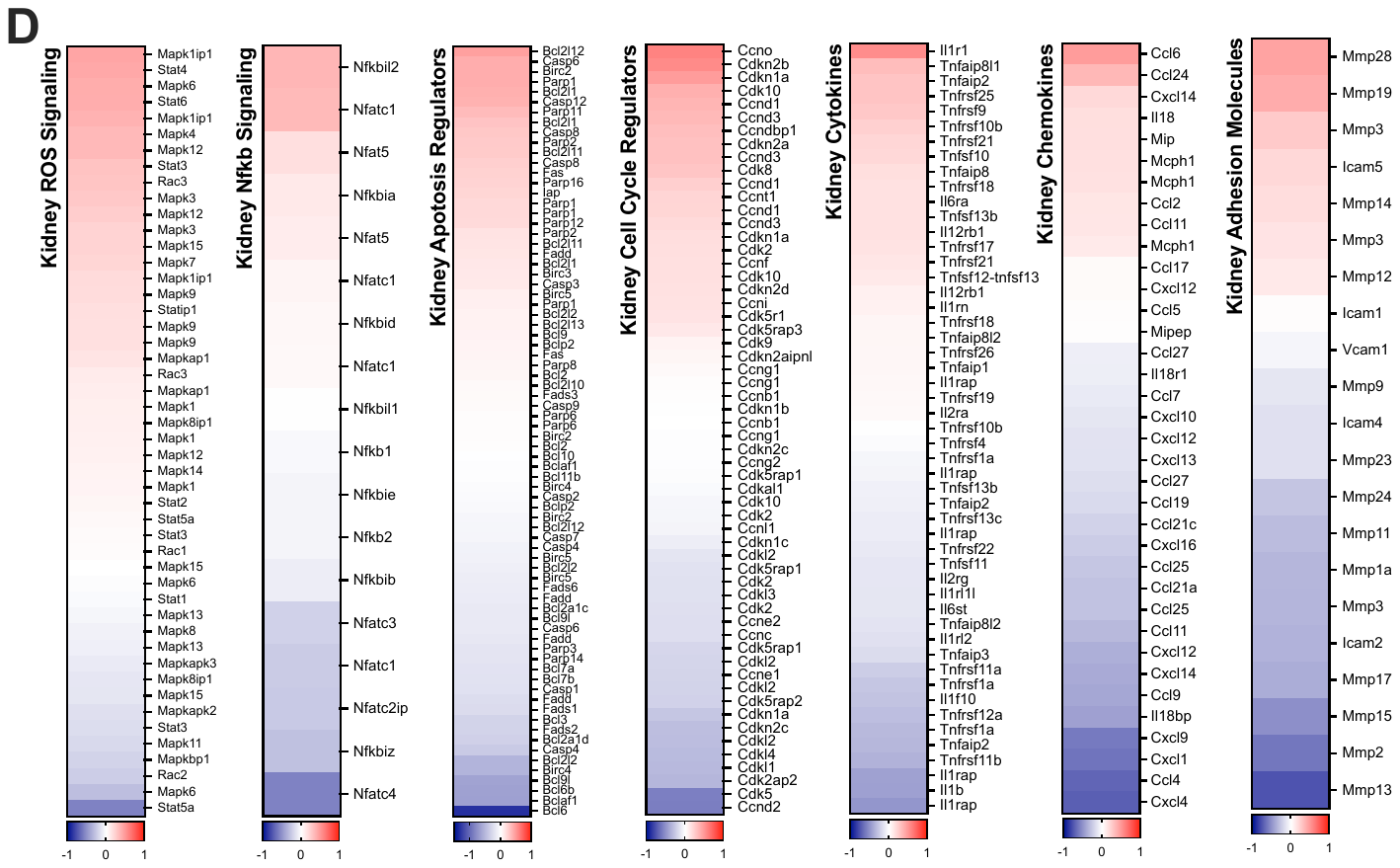
